# Lid Restriction Underlies Early nAngptl4-Mediated Lipoprotein Lipase Inhibition and Is Prevented by nAngptl4^E40K^ and Peptide Mimetics

**DOI:** 10.64898/2026.11.23.768672

**Authors:** Emma E. Lietzke, Mary S. Rouse, Ana F. Costa, Robb Welty, John T Melchior, Kayla G. Sprenger, Kimberley D. Bruce

## Abstract

The N-terminal fragment of Angiopoietin-like 4 (nAngptl4) potently inhibits lipoprotein lipase (LPL), the rate-limiting enzyme in triglyceride-rich lipoprotein (TRL) metabolism. However, the structural basis of nAngptl4-mediated inhibition remains poorly understood. Here, we present the first molecular dynamics (MD) simulations of nAngptl4, both in isolation and in complex with LPL. Our simulations reveal that nAngptl4 initiates LPL inhibition by restricting lid flexibility, rather than directly occluding the lipid pore, suggesting a reversible mechanism of inhibition that precedes longer-timescale destabilization. Simulations of the clinically relevant nAngptl4^E40K^ loss-of-function variant reveal that E40K does not grossly destabilize the coiled-coil architecture but instead redistributes local electrostatic interactions to alter predicted self-association interfaces, which may prevent oligomerization. Surprisingly, experimental studies demonstrate that nAngptl4^E40K^-derived peptides capturing the LPL-binding sequence increase LPL activity and attenuate nAngptl4-mediated inhibition by disrupting the interaction between full-length nAngptl4 and LPL. Together, these findings support a biphasic model in which early reversible inhibition is driven by lid restriction, followed by further LPL engagement and nAngptl4 polymerization, which is prevented by E40K. This work provides the first atomistic insight into nAngptl4-mediated LPL regulation and highlights opportunities for peptide-based therapeutic strategies that block nAngptl4-mediated LPL inhibition to improve TRL metabolism.

## Introduction

Lipoprotein lipase (LPL) is a central regulator of lipoprotein metabolism and lipid distribution across metabolically active tissues, including the brain^1–3^, heart^4^, muscle^5,6^, and adipose tissue^7^. Its primary function is to hydrolyze triglycerides contained within circulating triglyceride-rich lipoproteins (TRLs), thereby releasing free fatty acids for uptake and utilization by surrounding tissues^8–11^. Disruption of LPL activity through loss-of-function mutations, or elevated levels of endogenous inhibitors, has been linked to multiple disease states, including cardiovascular disease^12^, familial hypertriglyceridemia^13^, and Alzheimer’s disease^14^, highlighting its importance as a therapeutic target.

A major challenge to therapeutic development is the emerging complexity of LPL’s structure and function. Recent studies suggest that LPL can operate as an active monomer^15,16^ while also adopting multiple dimeric^16–19^ and higher-order^20,21^ assemblies. In addition, the enzyme exhibits a dynamic lid region^22–24^ and may undergo destabilization through inhibitor-induced perturbations in its N-terminal domain (NTD)^23,25^, particularly in regions surrounding the active site^26^ and lipid pore^18^. Together, these findings highlight the need for a more nuanced model of LPL structure–function relationships in which the most dynamic regions play a central role.

Angiopoietin-like 4 (Angptl4) has recently been identified as an inhibitor of LPL that destabilizes its NTD in both monomeric^23^ and dimeric forms^25,27–30^. Angptl4 is a soluble 406-residue protein expressed across multiple tissues and cell types, including adipocytes^31,32^, vascular endothelial cells^33^, and macrophages^34,35^. In a tissue-dependent manner^36^, Angptl4 can exist in both full-length and cleaved forms. Proteolytic processing by proprotein convertases^37,38^, such as furin, generates an N-terminal fragment^36,38,39^ (nAngptl4) and a fibrinogen-like C-terminal domain through cleavage at a central linker region (**Fig. 1A–B**). These domains serve distinct roles in lipid metabolism^40^, with nAngptl4 acting as a potent inhibitor of LPL^31,41,42^. Despite its well-established inhibitory function, the structural basis of LPL inhibition by nAngptl4 remains poorly understood.

**Figure 1.**
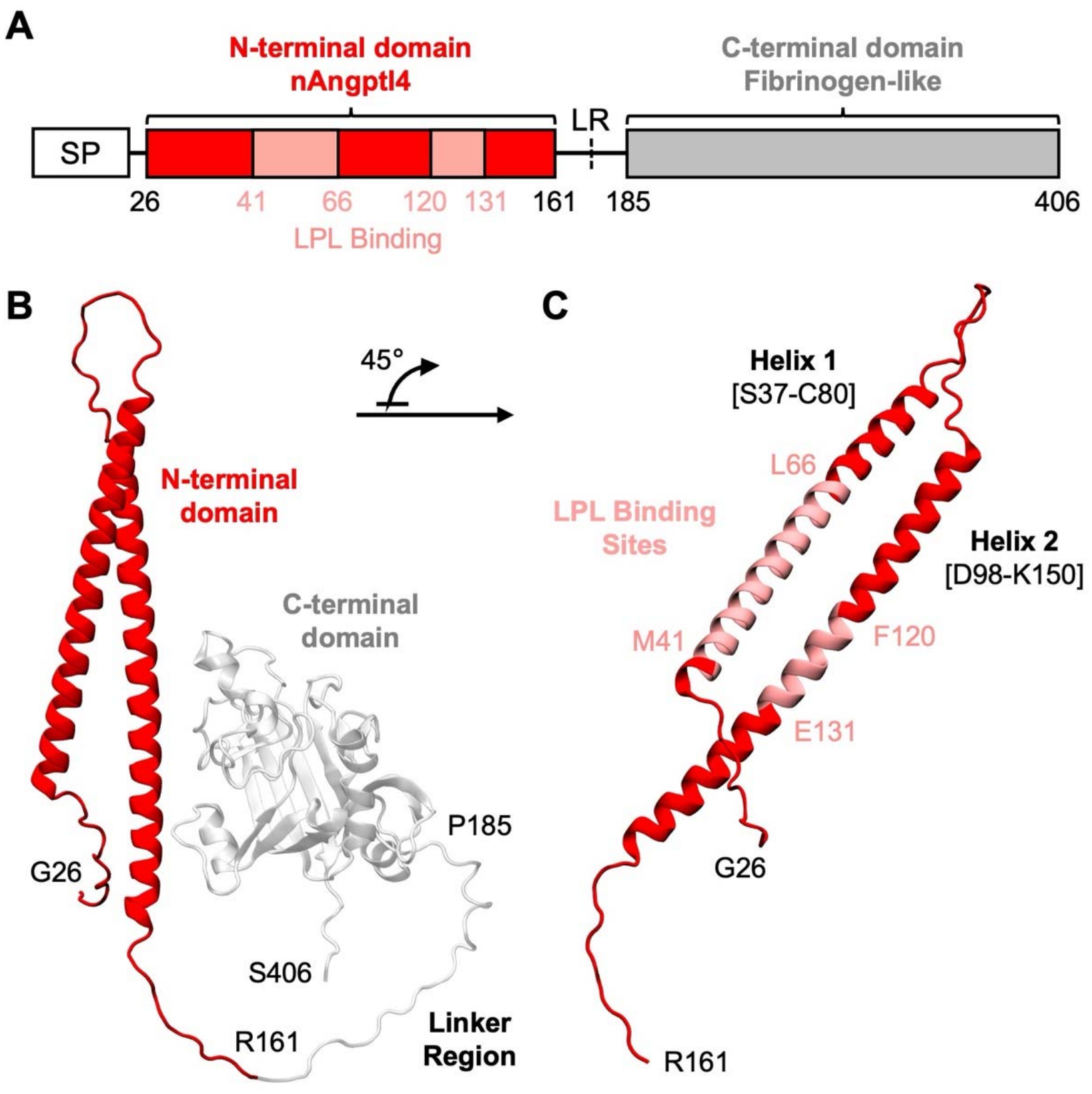
nAngptl4 contains multiple potential LPL-binding sites. (A) Schematic of Angptl4 domain organization. Angptl4 contains two major domains: the N-terminal domain (nAngptl4) and a C-terminal fibrinogen-like domain, connected by a linker region (LR) where the protein is cleaved extracellularly. The nAngptl4 domain contains multiple potential LPL-binding sites (pink). SP, signal peptide. (B) Full-length Angptl4 AlphaFold structure shown in cartoon representation. nAngptl4 is shown in red, while the LR and C-terminal are shown in white. (C) Structure of nAngptl4 (red) with key residues labeled and LPL-binding sites shown in pink. nAngptl4 adopts a coiled-coil structure formed by intertwined helices 1 and 2.

Early studies identified nAngptl4’s coiled-coil architecture^31,32,42,43^ and oligomerization behavior^41,43–45^, yet the precise mechanisms by which nAngptl4’s conformational state drives LPL inhibition remain unclear. In particular, the processes underlying nAngptl4 oligomer formation and how oligomerization alters its binding interfaces with LPL are not well resolved. This gap is likely due in part to the current lack of experimental structures of nAngptl4. Nevertheless, recent studies have begun to identify residues in both nAngptl4 and LPL that may mediate their interaction^46,47^ and contribute to LPL destabilization^23,25^ (**Fig. 1C**). However, these findings have not been examined in the context of the increasing complexity of LPL function and regulation. While molecular dynamics (MD) simulations could help resolve this complexity and provide critical insight into the interaction between nAngptl4 and LPL, nAngptl4 has not yet been studied using this approach.

One strategy for probing these interactions is to examine loss-of-function variants, such as Angptl4^E40K^, the most common variant of Angptl4, which is linked to reduced plasma TGs and a lower risk of both coronary artery disease^48–50^ and type 2 diabetes^51^. It is thought that the E40K mutation disrupts the formation of stable nAngptl4 oligomers, which are necessary for proper LPL inhibition^44^. However, the molecular basis for its reduced inhibitory activity toward LPL remains unclear. Resolving these mechanisms is essential for understanding both normal and impaired regulation of LPL, especially in the context of therapeutic development^52,53^. This need is further underscored by growing interest in other members of the angiopoietin-like protein family^54,55^, including Angptl3 and Angptl8. While Angptl3 shares structural similarities with Angptl4, it is a comparatively weaker inhibitor of LPL^30^ and requires formation of a 3:1 complex with Angptl8 for activity^56–58^. Meanwhile, Angptl8 also interacts with Angptl4 in a 1:1 manner that dampens the ability of Angptl4 to inhibit LPL independently^58,59^, further complicating our understanding of precisely how this protein family regulates LPL activity. Therapeutic strategies targeting the Angptl3/8 complex to modulate LPL activity have shown promise in alleviating dyslipidemia^60^, emphasizing the importance of defining the structure–function relationships that govern nAngptl4’s interactions with LPL.

To address these outstanding questions, we performed the first MD simulations of nAngptl4, both in isolation and in complex with LPL. These simulations were initiated from a complete structural model of LPL with a defined lid orientation, consistent with prior work^61^, to ensure a physiologically relevant starting point for assessing LPL lid and lipid pore dynamics. Our results show that nAngptl4 inhibits LPL primarily by restricting lid dynamics, suggesting a reversible mechanism of inhibition that precedes longer-timescale destabilization. Using a computational peptide-mapping approach, we identified key binding regions on nAngptl4 that interact with LPL, consistent with experimental observations. We further demonstrate that Angptl4-derived peptides function opposite to full-length nAngptl4, increasing LPL activity while also reducing nAngptl4-mediated inhibition, most potently in a peptide with the E40K substitution. Simulations of both full-length wild-type (WT) nAngptl4 and the E40K variant reveal that the monomers remain structurally stable but differ in their interaction networks with LPL centered on residue 40. While the E40K variant is largely restricted to self-interactions with a single neighboring residue, the WT form samples multiple transient salt bridges both internally and with LPL. These differences suggest a potential mechanism underlying the reduced function of E40K, namely weakened interactions with LPL and altered oligomerization behavior. Together, these findings provide new mechanistic insight into how nAngptl4 regulates LPL activity and highlight opportunities for targeting these interactions in cardiovascular disease.

## Methods

### Structural Model Preparation

The AlphaFold^62^ structure of human full-length Angptl4 was retrieved from the AlphaFold Protein Structure Database with the code AF-Q9BY76-F1-v4^63^. The Angptl4 AlphaFold structure comprises the entire protein domain (residues M1–S406; numbered 1–406). However, this structure contains the signal peptide domain, linker region, and Fibrinogen-like domain, which are cleaved prior to interacting with LPL. Thus, these regions (M1–G25 and R162–S406) were removed from nAngptl4. For the single-coil nAngptl4-derived peptides, the nAngptl4 structure was clipped to only contain the peptide portions (A36–L66 for H1, I116–L146 for H2). To create a single-coil H1-E40K peptide, PyMOL^64^ was used to mutate E40K in the pre-simulation H1 structure. For the H1X2 peptide, the structure was created by linking the H1 and H2 structures with a GGGGS linker sequence (e.g., H1-GGGGS-H2). The web server version of MODELLER^65^ was used to create the H1X2 structure, in which the full sequence was input and output a homology-modelled structure. The structure was refined in VMD^66^ prior to simulation to ensure proper steric alignment. For LPL–nAngptl4 simulations, the structure of LPL was taken from our prior work^61^. Specifically, the LPL-O structure was utilized. Residue numbering throughout refers to the full-length LPL and Angptl4 sequences including the signal peptide (residues 1–475 and 1–406, respectively); the simulated structures correspond to residues 28–475 (LPL) and residues 26–161 (full-length nAngptl4). Names of simulated Angptl4-dervied structures and their residue numbers are listed in **Table 1**.

**Table 1:** Angptl4-derived structures for simulations and corresponding residue numbers.

| **Structure Name** | **Amino acid [aa] Residue Numbers. + Modifications** |
|---|---|
| nAngptl4<sup>WT</sup> (WT) | 26–161 |
| nAngptl4<sup>E40K</sup> (E40K) | 26–161, with mutation E40K |
| H1 | 36–66 |
| H1-E40K | 36–66, with mutation E40K |
| H2 | 116–146 |
| H1X2 | 36–66:GGGGS:116–146 |

### Pre-Docking MD Simulations

All molecular dynamics simulations were carried out using GROMACS 2023 software^67^. Proteins and ions were represented with the CHARMM36 force field^68^, and water molecules were modeled with the TIP3P-modified forcefield^69^. Each system first underwent energy minimization in vacuum, followed by a second minimization step after solvation and ion neutralization to ensure structural relaxation. We then equilibrated the systems under NVT conditions for 1 ns at 310 K using a modified Berendsen thermostat^70^. This was followed by a 1 ns NPT equilibration phase at 310 K and 1 bar, employing the same thermostat together with the Berendsen barostat^70^. Production simulations of nAngplt4^WT^, nAngptl4^E40K^, and H1X2 were run independently in triplicate for 1 µs each, and production simulations of H1, H1-E40K, and H2 were run independently in triplicate for 2 µs each (replicate simulations denoted by Rep. 1–3). The total production time across all Angptl4 forms and replicates was 27 µs. All production simulation trajectories were generated using the Berendsen thermostat^70^ and the Parrinello– Rahman barostat^71^ maintained at 310 K and 1 bar, respectively. A 2-fs timestep was used, with full periodic boundary conditions and a 1.4-nm neighbor list cutoff. Bond constraints involving hydrogen atoms were enforced with the LINCS algorithm^72^. Long-range electrostatics were computed with the particle mesh Ewald (PME) method^73^ using a 1.2-nm real-space cutoff, and van der Waals interactions were cutoff at 1.2 nm.

### Selection of Structures for Docking

To evaluate structural stability and convergence during the production simulations, the root-mean-square deviation (RMSD) of each protein’s Cα atoms was calculated over time, using the corresponding energy-minimized structure as the reference. We also computed the root-mean-square fluctuation (RMSF) of the Cα atoms for each protein, averaging these values over the last 100 ns of each simulation. As with the RMSD analysis, the energy-minimized structure served as the reference for all RMSF calculations.

For all Angptl4 structures (full-length and peptide forms), the GROMACS clustering algorithm^67,74^ was utilized to identify the most representative structure across the three replicates, using the last 100 ns of each simulation. These structures (one per Angptl4 form) were selected for docking analyses with LPL. For H1X2 only, Rep. 2–3 simulations were used for clustering since the goal was to find a maintained paired-helix structure, and Rep. 1 lost this conformation.

### Protein–Protein Docking

For rigid protein–protein docking, the online docking server LZerD^75^ was employed. Six separate docking simulations were conducted for the following structures: nAngptl4^WT^, nAngptl4^E40K^, H1, H1-E40K, H2, H1X2, each paired with LPL-O structure from our previous work^61^. For each docked pair, 500 models were generated.

For nAngptl4–LPL docked models, models were assessed based on known binding sites between nAngptl4 and LPL, which had been previously determined via hydrogen–deuterium exchange/mass spectrometry (HDX-MS) LPL-Angptl4 experiments^47^ (e.g., aa41–66 on nAngptl4; aa76–87, aa109–126, aa245–251 on LPL). Distances between all atoms (excluding Hydrogen) on each protein were calculated with the distance module in the MDAnalysis library^76^ to determine contacts (cutoff < 5.0 Å) and hard clashes (cutoff < 2.0 Å). To assess the docked complexes, the percentage of expected LPL and nAngptl4 binding sites in contact, the number of total contacts, percentage of off-target contacts at expected binding sites, and the number of hard clashes were considered. The top-scoring model, based on these metrics, was determined and used for simulations. In VMD^66^, the top-scoring model, per respective complex, was inspected to confirm there were not any docking artifacts, such as unreasonable steric clashes.

For peptide–LPL docked models, models were not assessed with known binding sites considering that the interacting residues between the peptide and LPL would likely not be aligned exactly with those of full-length nAngptl4. Thus, the top-scoring complex from the LZerD^75^ ranking was used for simulations. These models were also assessed in VMD^66^ to ensure there were not any docking artifacts that would be biologically irrelevant.

For homodimeric nAngptl4 models, blind docking was also performed in LZerD^75^. To create trimer models, the top-scoring dimeric model (ranked by LZerD^75^) was docked to the representative monomeric structure, to produce homotrimeric models. To create tetrameric models, the top-scoring dimeric model was docked to itself, producing homotetrameric models. Since the exact interaction sites within the dimer interface are not known, preliminary analyses used the top 50 docked models, ranked on LZerD^75^, for contact maps. Top 500 docked models were used for solvent-accessible surface area (SASA) analyses.

### Post-Docking MD Simulations

Each Angplt4–LPL model was subjected to a 1-µs molecular dynamics simulation run, in triplicate, adding 18 µs of production sampling (45 µs of simulations in total). The docked complexes were simulated using the same workflow and parameters outlined above for the pre-docking MD simulations.

### Angptl4-Specific Simulation Trajectory Analysis

Model evaluation involved examining structural changes in Angptl4 across the simulations. Standard GROMACS tools were used to compute the Cα RMSD and RMSF for Angptl4 forms. Reference structures were the energy-minimized structures. For RMSF, the calculation was performed using the trajectory from the last 100 ns of each simulation. The area under the curve (AUC) was determined for different regions of the protein to identify significant changes in RMSF.

For the Angptl4 peptides, radius of gyration of the peptides was calculated using the radius_of_gyration module in MDAnalysis^76^. The distance between the terminal NTD and CTD Cα residues was calculated using MDAnalysis^76^ and the lingalg.norm module in Numpy^77^.

For full-length (FL) nAngptl4, a number of inter-residue calculations were performed. Salt bridge occupancies with residue 40 were calculated using the MDAnalysis library^76^. Qualifying distances between charged atoms, in charged residues (Asp, Glu, Lys, Arg; His excluded since it was□-protonated), with charged atoms in residue 40 were determined (cutoff < 4.0 Å) over the last 100 ns/simulation. Based on the charge of aa40, oppositely charged residues were filtered accordingly. Occupancy fractions of qualifying distances, per atom, were calculated. Similarly, interactions between hydrophobic residues in the core, from Helix 1 to Helix 2, were determined. To determine which residues would contribute, inwards-facing hydrophobic residues (Ala, Phe, Gly, Ile, Leu, Met, Pro, Val, Trp) were visually identified across the helices. Once determined, the heavy atoms in the sidechains of these residues were used for minimum distance calculations. The contact frequency per residue was determined for both WT and E40K, and the difference [WT-E40K] was calculated. To quantify potential bridging between the Helix 1 and Helix 2 within the full-length monomer, the number of H-bonds between these two regions (aa38–74 for Helix 1; aa99–148 for Helix 2) were calculated using the GROMACS^67^ hbond algorithm (bond distance cutoff = 0.35 nm; bond angle cutoff = 30°) over entire simulation time.

To determine all inter-residue contacts across full-length nAngptl4 models, the minimum distances between all heavy atoms per residue, to every other residue, were calculated using the MDAnalysis library^76^. Contact frequencies, based on these distances (cutoff < 5.0 Å), were determined over the last 100 ns of each simulation. The mean contact frequency per residue-residue, per variant, was determined. Differences in frequency [WT-E40K] were plotted.

Secondary structure assignments of Angptl4 residues were computed using the Compute DSSP module from the MDTraj Python library^78^. Calculations were performed over the entire simulation time and plotted against time. Assignments were listed as α-helix (H), residue in isolated β-bridge (B), extended strand participating in β-ladder, 3/10-helix (G), π-helix (I), hydrogen-bonded turn (T), bend (S), and loop/irregular elements (C). For nAngptl4, the fractional α-helicity of residues across the entire protein, over the last 100 ns, was calculated.

To test for significant differences across Angptl4 variants, a Welch’s t-test was used (n = 3 simulations) to compare the means. A Hedges’ g test was used to measure effect size, considering the small sample size of simulations.

### Angptl4–LPL Simulation Trajectory Analysis

Cα RMSD for the Angptl4–LPL complex, LPL alone, and Angptl4 alone were calculated in the same manner as described above. RMSF for Angptl4 forms and LPL were calculated as described above, using the last 100 ns of the trajectory. Reference structures were the energy-minimized structures.

The SASA of the LPL residues in the lid domain and residues surrounding the lipid pore (aa245– 260; aa81–91, 114–128, 159–161, 193–192, 210–223, 261–269, respectively; LPL numbered 1–475) were calculated using the shrake_rupley module in the MDTraj Python library^78^, from 900– 1000 ns in each simulation. Here, SASA measured the surface area of all atoms exposed to solvent, using water, lipid, and protein as the probe sizes (radii = 0.14, 0.50, and 1.00 nm, respectively). SASA values of LPL, in the presence of a ligand, were normalized against the SASA of ligand-free LPL, specific to each frame in each trajectory per simulation. Normalized SASA values ranged from 0–1, with 0 corresponding to no ligand coverage and 1 corresponding to complete ligand coverage, over regional residues and time. To test for significant differences across variants (peptide and full-length), a Welch’s t-test was used to compare the means.

Lid distance of LPL was calculated using the same methods as described previously^61^ (e.g., distance between the Cα P187–G256 with Python library MDAnalysis^76^). Delta lid distance was plotted over time, using the lid distance in the first frame of the production trajectory as the comparison point.

The LPL residues within 4.5 Å of Angptl4 (both peptides and full-length), over the last 100 ns of each simulation, were calculated using the distances module in the MDAnalysis Python library^76^. Interaction frequencies for LPL residues were normalized over the computed time and plotted. The LPL and nAngptl4 residues within 5 Å of each other^79^, over the last 100 ns of each simulation, were calculated using the compute_distances module in the MDTraj Python library^78^. Distances were plotted in contact frequency maps per simulation complex normalized over time (e.g., full-length WT, full-length E40K). The mean contact frequency per residue-residue, per variant, was determined. Differences in frequency [WT-E40K] were also plotted. These residues were compared to those identified from spectroscopy assays in existing literature.

Binding interactions between LPL and Angptl4 were quantified by estimating interaction free energies using the molecular mechanics Poisson–Boltzmann surface area (MM/PBSA) method. Calculations were performed with the gmx_MMPBSA tool^80^ integrated within GROMACS^67^ and focused on the final 100 ns of each trajectory (900–1000 ns). To further resolve residue-level contributions, we carried out energy decomposition analyses to identify key contributors to binding and assess their interaction frequencies. For each LPL–Angptl4 complex, free energies were sampled at 0.5-ns intervals over the analysis window, resulting in 201 measurements per simulation. Energy distributions were visualized as boxplots with associated standard deviations, and statistical comparisons between groups were performed using Welch’s t-test. Decomposition results are presented for the top 20 stabilizing residues in LPL, the top 20 residues in full-length nAngptl4, and the top 5 residues in the nAngptl4-derived peptides.

### nAngptl4 Oligomer Analysis

Homodimeric models of nAngptl4^WT^ and nAngptl4^E40K^ were generated (see **Protein–Protein Docking**). Contact maps between the monomers within the homodimer (called Monomer 1 and Monomer 2) were created using the MDAnalysis library^76^, across the top 50 docked models (see **Angptl4-Specific Simulation Trajectory Analysis** for more details on contact map calculation).

The solvent-accessible surface area (SASA) of homodimeric, homotrimeric, and homotetrameric components were calculated using the shrake_rupley module in the MDTraj Python library^78^, for the top 500 models for both variants. Here, SASA measured the surface area of all atoms exposed to solvent, using water as the probe size (radii = 0.14 nm). The SASA of residues 40–66 in all chains was calculated and compared across oligomers and variants. The SASA of the LPL lipid pore and lid coverage, with each variant’s dimer, trimer, and tetramer, was determined used the same method as described above (see **Angptl4–LPL Simulation Trajectory Analysis**). To test for significant differences across Angptl4 variants, a Welch’s t-test was used (n = 500 docked models) to compare the means.

### Simulation Trajectory and Analyses Visualization

All simulation trajectory images were created in VMD^66^. Computational-based plots were created in Python. Experimental-based plots were created in BioRender Graph (https://BioRender.com).

### Peptide Design and Synthesis

Custom peptides were synthesized by GenScript 95% purity or above. Lyophilized peptides were resuspended in 100mM sodium phosphate buffer (pH 7.4) immediately before use.

### Fluorescence Detection of Lipoprotein Lipase Activity at Elevated Temperature Using a Lipid-Free EnzChek^TM^ Lipase Substrate

Lipid-free LPL activity was quantified using an optimized EnzChek^TM^ assay^81^ as previously described^82^. Assays were performed in 96-well plates with ten conditions assayed in quadruplicate: vehicle control, LPL alone, nAngptl4 alone, LPL + N-terminal fragment (nAngptl4), LPL + H1, LPL + H1-E40K, LPL + H2, LPL + H1 + nAngptl4, LPL + H1-E40K + nAngptl4, and LPL + H2 + nAngptl4. Each Angptl4 peptide was added at 200 ng per well. Samples were first incubated without nAngptl4 for 7.5 minutes, after which 200 ng of nAngptl4 was added to the appropriate wells and incubated for an additional 7.5 minutes. Bovine LPL (diluted 1:2000) and Angptl4 samples were diluted in 100 mM sodium phosphate buffer (pH 7.4). An 80× EnzChek^TM^/Zwittergent working solution was prepared by mixing equal volumes of EnzChek^TM^ lipase substrate (green fluorescent, 505/515; Thermo Fisher Scientific, E33955) and Zwittergent 3 12 detergent (Sigma, 693015), yielding final concentrations of 50 µM substrate and 4% Zwittergent. Then, 50 µL of molecular-grade water and 47.5 µL of 4× sample buffer were added to each well. Immediately before measurement, 2.5 µL of the EnzChek^TM^/Zwittergent working solution was added to each well for a final substrate concentration of 0.625 µM in a 200 µL volume. Fluorescence was measured at 490/515 nm excitation/emission using a gain of 100, normal read speed, and a 7 mm read height. Readings were collected every 5 minutes for 1 hour at 37°C.

### Circular Dichroism Experiments

Circular dichroism (CD) spectroscopy was performed to determine the secondary structure of peptides by measuring the differential absorption of left-and right-circularly polarized light. CD quantifies the presence of α-helices, β-sheets, and random coils to determine peptide makeup. CD spectra were collected for human Angptl4 peptides at room temperature in the presence of the sodium phosphate buffer. Quantification of secondary structure from CD spectra was determined using the Beta Structure Selection (BeStSel) method^83^. The BeStSel server is freely accessible at https://bestsel.elte.hu.

## Results

### Full-length nAngptl4 binding to LPL’s lid domain restrains lid dynamics, suggesting a reversible inhibition mechanism at short timescales

nAngptl4 is a well-established inhibitor of LPL. Here, we recapitulate this effect, showing that nAngptl4 robustly suppresses LPL-mediated hydrolysis of a lipid analogue (**Fig. 2A**, **Supplementary Table 1**). nAngptl4 has long been thought to destabilize LPL, potentially by promoting dissociation of the LPL dimer^25,27–30^. However, recent advances in our understanding of LPL structure and function have uncovered additional complexity in its regulation, including evidence for active LPL monomers^15,16,23^, structurally variable dimers^16–19^, and more complex lid^24,61^ and lipid pore dynamics^18^. As such, the mechanisms by which nAngptl4 inhibits LPL are more complex than initially appreciated and require further characterization.

**Figure 2.**
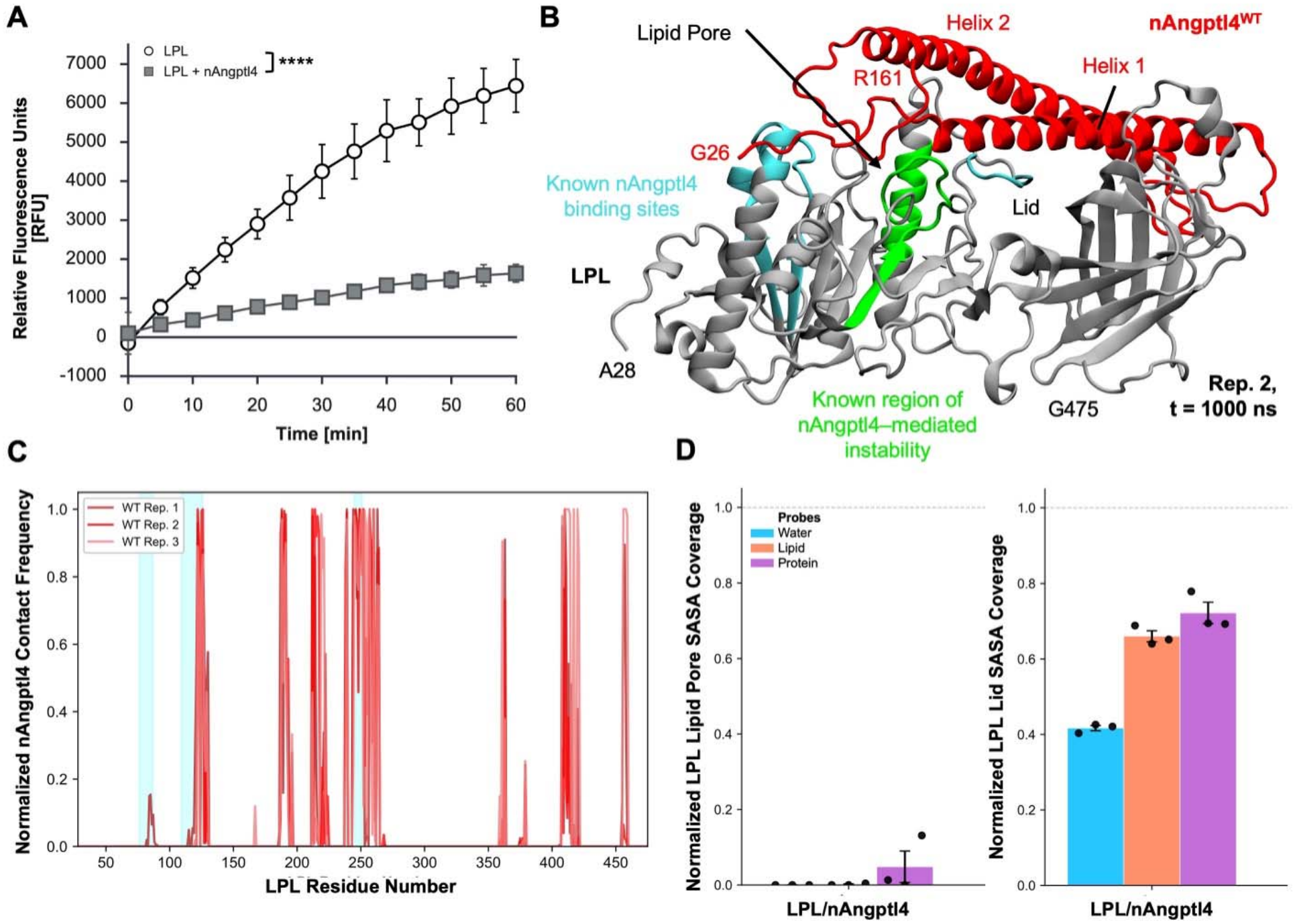
nAngptl4 interacts primarily with the lid of LPL to inhibit active site access. (A) nAngptl4 reduces LPL-mediated hydrolysis of a fluorescent triglyceride analog. **** = P<0.001, differences between groups were analyzed by two-way ANOVA with Tukey’s post hoc multiple-comparisons test (n=4); see Supplementary Table 1 for complete statistics. (B) Front view of LPL bound to nAngptl4^WT^ (final simulation time point, t = 1000 ns, for Replicate (Rep.) 2). LPL is shown in gray, with prior identified nAngptl4 binding sites in cyan (amino acids (aa) 76– 87, 109–126, 245–251; LPL numbered aa 1–475) and regions previously shown to undergo allosteric instability in response to nAngplt4^WT^ binding in green (aa 205–220, 264–274). nAngptl4^WT^ is shown in red. (C) Normalized contact frequency of nAngptl4^WT^ across LPL residues from 900-1000 ns. Contact values range from 0 to 1, where 0 indicates no contact and 1 indicates persistent contact. Cyan regions denote nAngptl4 binding sites on LPL. (D) Normalized solvent-accessible surface area (SASA) coverage of nAngptl4^WT^ on LPL for the lipid pore entrance-surrounding residues (left) and lid residues (right) from 900–1000 ns. Coverage is normalized to unbound LPL and ranges from 0 to 1, where 0 indicates no coverage, and 1 indicates complete coverage. Probe radii correspond to water, lipid, and protein (0.14, 0.50, and 1.00 nm, respectively). Mean coverage values are shown with standard error of the mean (SEM) by black bars.

In our previous work^61^, we proposed that lid flexibility is an important regulator of LPL activity and found that it can be enhanced by a peptide mimetic of the known LPL activator apolipoprotein C-II (ApoC-II^61,82^). We therefore hypothesized that the opposite occurs during inhibition—that loss of lid flexibility (i.e., lid stabilization) contributes to LPL inactivation and eventual destabilization. To test this hypothesis, we performed MD simulations of nAngptl4– LPL interactions. nAngptl4 and LPL were first simulated independently in triplicate (1000 ns each), followed by rigid docking to generate complex models. Models were evaluated based on agreement with experimentally identified interaction residues, and the top-scoring complex was subsequently simulated in triplicate (1000 ns each).

Monomeric nAngptl4 remained stably bound to LPL in all simulations, exhibiting low root-mean-square deviation (RMSD) fluctuations (**Supplementary Fig. 1A**) and minimal reorientation relative to the initial docked pose (**Supplementary Fig. 1B–C**). Notably, nAngptl4 localized across the top of LPL, positioning over the lid and lipid pore (**Fig. 2B**). These results are consistent with HDX-MS data^46,47^, which show stable interactions of nAngptl4 with residues surrounding the lipid pore in the N-terminal domain (NTD), lid region, and C-terminal domain (CTD) near the lipid-binding site (**Fig. 2C**; **Supplementary Fig. 1D**). While interactions within the NTD were expected, the observed bridging of nAngptl4 across the NTD and CTD suggests a potential mechanism for reversible inhibition^45^ or interference with LPL dimerization^25,27–30^. In addition, these findings support the ability of our simulations to capture biologically relevant interactions between nAngptl4 and LPL.

We next asked whether nAngptl4 inhibits LPL through a two-step mechanism: first, by restraining lid flexibility and sterically blocking access to the lipid pore, and second, by inducing longer-timescale destabilization of the NTD. Although our simulations did not capture this latter process, we instead evaluated the short-timescale effects of nAngptl4 binding on LPL structure and dynamics. Solvent-accessible surface area (SASA) analysis of the lid and lipid pore regions (**Supplementary Fig. 1E**) revealed that access to the lipid pore was largely unchanged in the presence of nAngptl4 (**Fig. 2D**, left). In contrast, nAngptl4 significantly reduced lid accessibility, as evidenced by increased SASA coverage of the lid region (**Fig. 2D**, right), which corresponded to decreased lid dynamics (**Supplementary Fig. 1F**). These results suggest that inhibition is initially driven by strong lid binding that restricts lid flexibility. Consistent with this, residues from both the primary^43,46,47,84^ and secondary^46^ previously defined LPL-binding regions of nAngptl4 (**Fig. 1A, C**) consistently engaged the lid throughout the simulations (**Supplementary Fig. 1D**). Since experimental data identified these same regions (Add ref), we next sought to determine which residues within them contribute most strongly to LPL binding using a computational peptide-mapping approach.

### nAngptl4-derived peptides attenuate nAngptl4-mediated LPL inhibition by blocking lid engagement

Building on our previous work identifying a peptide mimetic of the LPL activator ApoC-II^82^, we asked whether a similar strategy could achieve the opposite effect—LPL inhibition—by isolating LPL-binding regions within nAngptl4. Of the two LPL-binding regions identified in previous studies^46,47^, the Helix 1 region is generally recognized as the primary LPL-binding site^43,46,47,84^ of nAngptl4 (**Fig. 1A, C**). Notably, this site lies adjacent to the E40K point mutation, the most common Angptl4 variant^48–51^, with reduced ability to inhibit LPL.

To probe these interactions at the molecular level, we designed four peptides comprising these regions. For the primary LPL-binding site, we designed peptides H1 (A36–L66) and H1-E40K (A36–L66 containing the E40K mutation). For the secondary binding site, we designed peptide H2 (I116–L146). Because these peptides represent isolated helical segments, which are likely both required for full lid engagement and LPL inhibition, we also designed a fourth construct, H1X2, which bridges the two regions creating a short mimetic of nAngptl4’s coiled-coil domain using a 5-residue linker (GGGGS) between H1 and H2. All peptides were simulated in triplicate until their secondary structures converged, with 2000-ns simulations for the single-coil peptides (H1, H1-E40K, H2) and 1000-ns simulations for H1X2.

The single-coil peptides were largely unstable over the course of the simulations, as reflected in fluctuating RMSD profiles (**Supplementary Fig. 2A**), whereas H1X2 converged rapidly in two of three replicates. Differences in peptide stability were further supported by radius of gyration and NTD–CTD distance measurements (**Fig. 3A**; **Supplementary Fig. 2B**), which showed that coiled-coil formation stabilizes peptide compactness and inter-terminal distance. Final trajectory snapshots showed substantial loss of α-helicity in the single-coil peptides relative to the corresponding regions in full-length nAngptl4 (**Supplementary Fig. 2C–F**). Specifically, H1 and H2 retained small helical segments, whereas H1-E40K lost nearly all structure. In contrast, H1X2 maintained α-helicity, even in the partially folded replicate (Rep. 1), and showed near-complete structural preservation in the other replicates (Reps. 2–3). Secondary structure analysis using the Dictionary of Secondary Structure in Proteins (DSSP) method confirmed these trends, showing rapid loss of α-helicity in the single-coil peptides, most prominently for H1-E40K (**Supplementary Figs. 2–3**). These differences are reflected in representative post-simulation structures (**Fig. 3B**).

**Figure 3.**
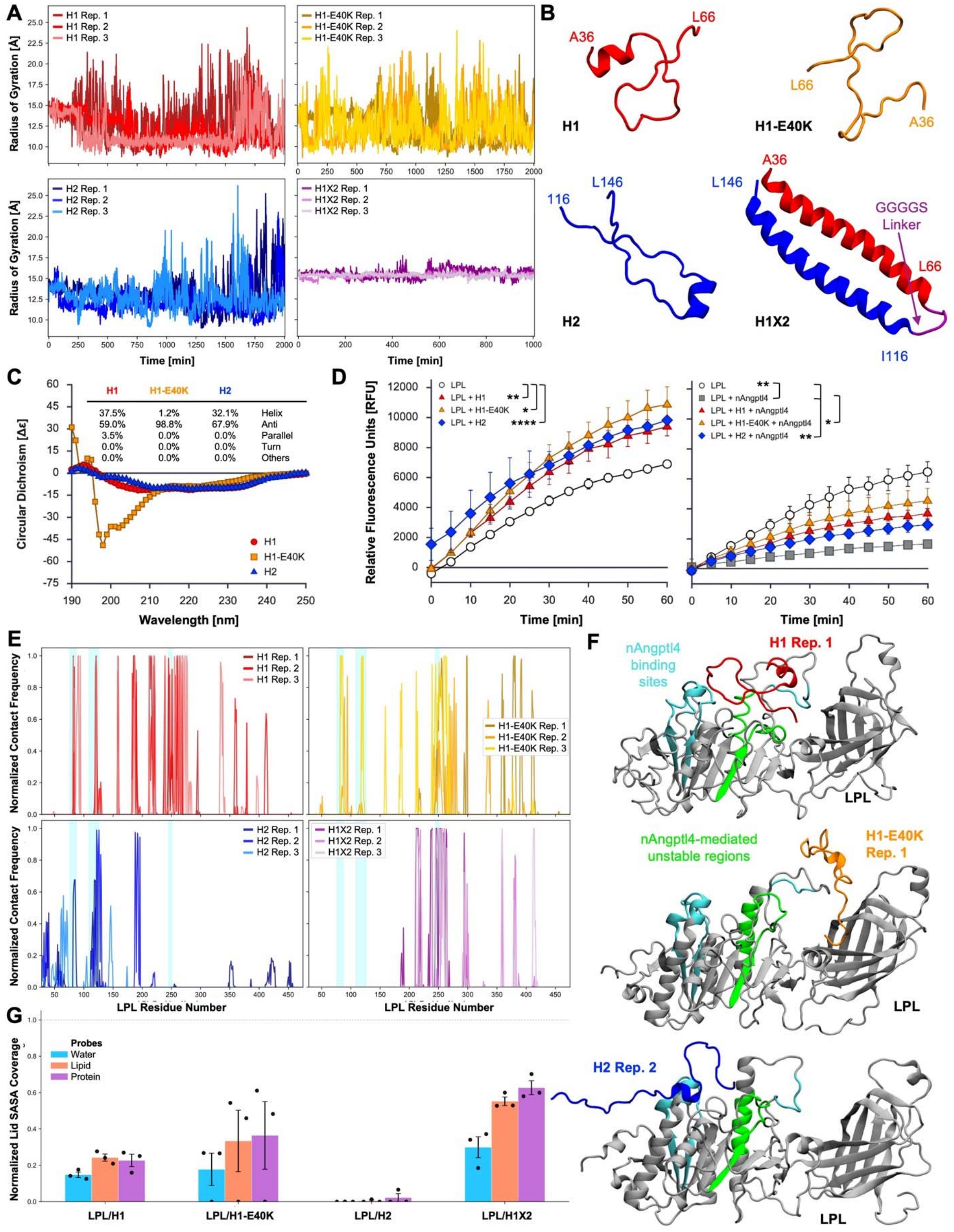
Peptide H1 suggests nAngptl4’s primary binding site for interactions with LPL. (A) Radius of gyration of nAngptl4-derived peptides over the course of the simulations. H1 is shown in shades of red, H1-E40K in orange, H2 in blue, and H1X2 in purple. (B) Representative post-simulation structures of H1 (red), H1-E40K (orange), H2 (blue), and H1X2 (red, purple, blue). (C) Circular dichroism spectra for H1, H1-E40K, and H2 in sodium phosphate buffer. Secondary structure was estimated using Beta Structure Selection (BeStSel), where “Helix” corresponds to α-helix, “Parallel” to parallel β-sheet, and “Anti” to anti-parallel β-sheet or lack of secondary structure. (D) H1, H1-E40K, and H2 activate LPL and inhibit nAngptl4-mediated inhibition of LPL (n = 4, except for ‘LPL + H1 + nAngptl4’ where n = 3) * = P<0.05, **= P<0.01, *** P<0.001, differences between groups were analyzed by two-way ANOVA with Tukey’s post hoc multiple-comparisons test (n=4); see Supplementary Tables 2– 3 for full statistics. (E) Normalized contact frequency of peptides across LPL residues from 900–1000 ns. Cyan regions denote nAngptl4 binding sites on LPL. (F) Front view of LPL bound to peptides (final simulation time point, t = 1000 ns, for representative H1, H1-E40K, and H2 replicates (Rep.)). LPL is shown in gray, with nAngptl4 binding sites in cyan and allosterically unstable regions in green. (G) Normalized SASA coverage of peptides on LPL for lid residues from 900–1000 ns. All normalization procedures, residue ranges, probe definitions, and analysis methods are as described in Figure 2.

The loss of α-helicity in peptides H1, H1-E40K, and H2 was confirmed by circular dichroism (**Fig. 3C**). Given the practical challenges associated with synthesizing H1X2, this construct remains purely theoretical. We found that H1 and H2 retained some helicity while H1-E40K was mostly disordered, supported by the very negative signal at 195 nm^85^. These trends are consistent with the DSSP profiles near the end of the simulations (**Supplementary Figs. 3C, 4C**), suggesting that the E40K mutation shifts the peptide away from its canonical α-helicity. However, the DSSP profiles and circular dichroism data differ in their predictions of β-sheet and irregular characters, likely an artifact from solution differences in the experimental sodium phosphate buffer versus the computational 150 mM NaCl solution.

Surprisingly, H1, H1-E40K, and H2 all increased LPL activity when measuring hydrolysis of a lipid analogue (**Fig. 3D**, left; **Supplementary Table 2**). This behavior was unexpected, considering the potent inhibitory nature of endogenous, full-length nAngptl4. Given the reduced α-helical content of the nAngptl4-derived peptides, it is plausible that their inability to inhibit LPL stems from disruption of the secondary structure required for nAngptl4-mediated inhibition.

Since H1, H1-E40K, and H2 partially encompass the LPL-binding region of full-length nAngptl4, we reasoned that these peptides may block the interaction between nAngptl4 and LPL, thereby preventing nAngptl4-mediated inhibition of LPL activity. We found that all three peptides significantly attenuated nAngptl4-mediated inhibition, although none fully reversed its inhibitory effect (**Fig. 3D**, right; **Supplementary Table 3**). Moreover, H1-E40K, which is also the most “unstructured” peptide, exhibited the strongest ability to both activate LPL and inhibit nAngptl4. While E40K is a known loss-of-function variant, its ability to counteract the inhibitory effects of nAngptl4^WT^ has not been previously described. Indeed, this striking dual-function behavior, whereby the peptides both increased LPL activity and attenuated nAngptl4-mediated inhibition, was unexpected.

To investigate the molecular basis of attenuated nAngptl4-mediated inhibition by H1, H1-E40K, and H2, representative peptide structures were docked to an open-lid conformation of LPL, in which the active site is accessible (see previous work^61^) and simulated in triplicate (1000 ns each). Despite unbiased (i.e., “blind”) docking, the peptides consistently clustered near the active site and lid of LPL, aligning with HDX-MS–identified binding regions for full-length nAngptl4 (**Supplementary Fig. 5A–B**). H1 and H1X2 were the most stably bound during subsequent MD simulations, exhibiting only minor reorientation. In contrast, H1-E40K showed increased fluctuations and dissociated in one simulation, and H2 bound weakly and only rarely engaged the lid (**Supplementary Fig. 5C–E**). Interaction frequency maps showed that H1 robustly engaged key LPL-binding regions, including the lipid pore, and lid region (residue ∼250) whereas H2 primarily engaged more distal regions (**Fig. 3E**). H1X2 interactions were largely localized to lid residues within the NTD, notably excluding other predicted binding sites that the single-coil peptides were able to engage (**Fig. 3E**). Unstructured H1-E40K exhibited a similar interaction profile, with a contact cluster around the lid-domain and reduced contact at the lipid pore (**Fig. 3E**). Visual analysis confirmed flexible positioning of the peptides across LPL (**Fig. 3F**). This behavior is reminiscent of unstructured ApoC-II-P which can bridge the N and C-terminal domains of LPL to simultaneously preserve overall enzyme structure while maintaining lid flexibility to preserve activity.

To compare mechanisms of inhibition, we quantified LPL lipid pore and lid coverage by the peptides through SASA calculations. Most peptides, except H2, exhibited greater lipid pore coverage than full-length nAngptl4 (**Supplementary Fig. 5F vs. Fig. 2D, left**). In addition, H1 and H2 were less effective at covering the lid. While H1-E40K and H1X2 exhibited more lid coverage, H1X2 showed coverage comparable to full-length nAngptl4 (**Fig. 3G vs. Fig. 2D, right**). These results suggest a trade-off between lipid pore and lid engagement for small peptides versus large, endogenous proteins, and indicate that the peptides may mediate a distinct activation mechanism that enables lid flexibility while also still interacting with the lipid pore (**Supplementary Figs. 1F, 5G–H**). Notably, LPL experienced the largest degree of lid flexibility in the presence of H1-E40K, aligning with our hypothesis that the promotion or restriction of lid dynamics correlates with LPL activation and inhibition, respectively. Collectively, these findings demonstrate that LPL activity can be modulated through multiple mechanisms by different regions of the same protein, enabling graded levels of activation with simultaneous inhibition of endogenous inhibitors.

*In silico* binding free energy calculations further supported the lipid pore and lid as key interaction hotspots. H1, H1-E40K, and H1X2 exhibited similar binding affinities for LPL, while H2 showed significantly weaker interactions (**Supplementary Fig. 7A**). Consistent with our earlier interaction frequency and LPL coverage analyses, these results confirm that residues within H1 constitute the primary LPL-binding site in nAngptl4. Residue-level contributions to binding aligned with regions identified by HDX-MS^47^ (**Supplementary Fig. 7B**; **Supplementary Fig. 8A–D**), and peptide residues R56, R61, and R63 consistently emerged as dominant energetic contributors (**Supplementary Fig. 8A–B**). Notably, residue 40 contributed to LPL binding in H1 but not in H1-E40K, suggesting altered interactions in the mutant. Together, these results support a model in which the E40K mutation impairs LPL inhibition by weakening binding interactions and potentially altering both LPL engagement and nAngptl4 self-association.

### E40K redistributes intramolecular contacts in full-length nAngptl4 while preserving its coiled-coil architecture

To determine whether the effects of variation at residue 40 observed in the nAngptl4-derived peptides are preserved in the full-length construct, we next simulated nAngptl4^E40K^ (amino acid (aa) 26–161) in explicit solvent for 1 µs in triplicate and compared the results with our earlier simulations of nAngptl4^WT^. Both variants exhibited an initial increase in RMSD before reaching a plateau, after which it remained stable for the duration of the simulations (**Supplementary Fig. 9A**). Visual inspection confirmed preservation of the paired helices in the coiled-coil domain, with flexibility concentrated in the hinge and terminal regions (**Supplementary Fig. 9B–C**).

Root-mean-square fluctuation (RMSF) profiles calculated over the final 100 ns supported the above observations (**Fig. 4A**). Specifically, statistically significant differences between the variants localized primarily to full-length Helix 2, with nAngptl4^WT^ exhibiting greater flexibility in its C-terminal region at residues spatially adjacent to residue 40 in full-length Helix 1 (**Fig. 4 A–B**; see aa124-148 shown in brackets). Our earlier analyses indicated that full-length Helix 2 does not constitute a primary interface for LPL binding; however, the increased flexibility observed in this region of nAngptl4^WT^ suggests that it may sample alternative conformations or transient interactions that could influence LPL engagement. In contrast, this flexibility is reduced in the E40K variant, indicating a more constrained local structural environment. These differences may contribute to changes in the overall interaction landscape of nAngptl4 in the presence of the E40K mutation.

**Figure 4.**
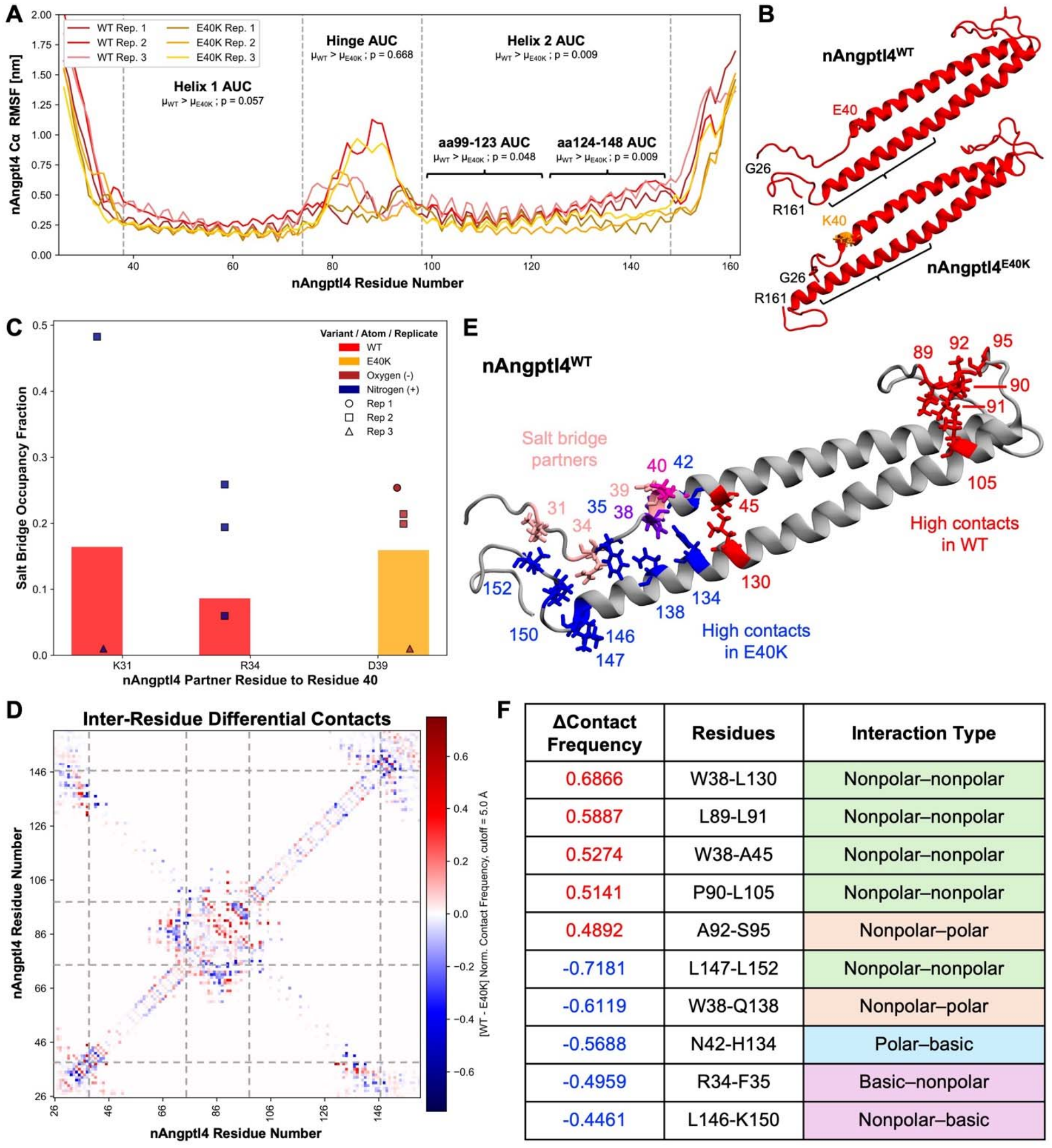
nAngptl4^E40K^ maintains the coiled-coil architecture but alters transient salt bridge formation. (A) Root-mean-square-fluctuation (RMSF) of the Cα backbone of nAngptl4^WT^ (red) and nAngptl4^E40K^ (orange) over the full simulation time. Defined regions include: Helix 1 (aa 38–74), hinge (aa 75–98), and Helix 2 (aa 99–147). Area-under-the-curve (AUC) of RMSF was calculated for each variant. Statistical significance between variant means was assessed via Welch’s t-test. (B) Representative post-simulation structures of nAngptl4^WT^ (top) and nAngptl4^E40K^ (bottom). Residue 40 is shown in licorice representation (red for nAngptl4^WT^, orange for nAngptl4^E40K^). Black brackets correspond to aa124–148. (C) Salt bridge occupancy for residues interacting with position 40 over the final 100 ns. Mean occupancies for nAngptl4^WT^ (red) and nAngptl4^E40K^ (orange) are shown as bars and individual contacts as points. (D) Differences in nAngptl4 inter-residue contact frequency over the final 100 ns. Positive (red) and negative (blue) values indicate higher contact frequency in nAngptl4^WT^ and nAngptl4^E40K^, respectively. (E) Representative post-simulation structure of nAngptl4^WT^ highlighting salt bridge partner residues (light pink), residue 40 (hot pink), and residues exhibiting differential contacts (red, purple, blue). (F) Top differential contacts for nAngptl4^WT^ (red) and nAngptl4^E40K^ (blue) (data derived from (D)).

To evaluate if these differences in flexibility reflected changes in secondary structure, DSSP analysis was performed over the simulation trajectories (**Supplementary Fig. 10A–B**). Both variants displayed modest reductions in α-helicity during the first 500 ns, consistent with early RMSD changes, but retained well-defined helices in representative late-stage structures (**Supplementary Fig. 9B–C**). Mean fractional α-helicity calculated over the final 100 ns was not statistically significant between variants, likely reflecting that the E40K variant is not broadly destabilizing nAngptl4 secondary structure in the timescales observed here (**Supplementary Fig. 10C**).

Because coiled-coil stability depends on inter-residue interaction networks^86^, we next examined whether the E40K substitution altered electrostatic contacts. Given the charge reversal at residue 40, we quantified salt bridge occupancies over the final 100 ns (**Fig. 4C, E**). In nAngptl4^WT^, E40 engaged transiently with residues K31 and R34 (occupancies: µ_31_ = 0.164 ± 0.276, µ_34_ = 0.086 ± 0.149), whereas K40 in the mutant formed interactions primarily with D39 (µ_39_ = 0.159 ± 0.131). Although occupancies were low in both variants, the broader interaction pattern in nAngptl4^WT^ suggests that the native glutamate at residue 40 participates in a more distributed electrostatic network in this region. Despite WT and E40K have similar overall flexibility in Helix 1 (**Fig. 4A**), Replicates 2 and 3 of the WT simulations exhibit a higher RMSF in ∼aa40–50, which coincides with both being involved in salt bridges (although minimally in Rep. 3). This may suggest that upstream salt bridge formation, particularly at residues not directly next to residue 40, could have allosteric effects that ultimately influence the rigidity of the LPL binding region.

To assess whether mutation-induced effects extended beyond local electrostatics, we compared residue–residue contact frequencies across the full-length N-terminal fragment (**Fig. 4D**). As expected, the largest differences between the two full-length variants occurred in intrinsically flexible regions, including the termini and hinge. Notably, the largest differences in contact frequency were enriched in the hinge for nAngptl4^WT^, whereas E40K showed relatively greater contributions from residues in the N-and C-terminal tail domains (**Fig. 4E**). In nAngptl4^WT^, these prominent contacts were largely nonpolar, while E40K displayed a broader mix of side chain chemistries (**Fig. 4F**). Despite these shifts, the hydrophobic core characteristic of the coiled-coil domain remained largely unchanged: nonpolar inward-facing helical contacts did not differ significantly between variants (**Supplementary Fig. 11A–C**), nor did inter-helical hydrogen bonding (**Supplementary Fig. 11D**).

Collectively, these data indicate that E40K does not induce large-scale disruption of the coiled-coil architecture. Instead, the mutation is associated with subtle redistribution of interaction networks within the protein, including reduced diversity of local salt bridges near residue 40 and shifts in contact enrichment between hinge and terminal regions. These changes coincide with reduced flexibility in full-length Helix 2 in the mutant, particularly in residues proximal in space to residue 40. These modest, spatially distributed effects are consistent with a small allosteric perturbation that could influence stability or binding over longer timescales than captured in these simulations. These findings support the hypothesis that E40K exerts a biphasic effect on nAngptl4 structural dynamics and LPL inhibition: (1) an early stage characterized by reduced affinity for LPL, and (2) a later stage in which allosteric perturbations accumulate in the unbound protein, impairing proper oligomerization. As a result, nAngptl4^E40K^ may become progressively less stable over time, with reduced oligomer formation ultimately diminishing its ability to inhibit LPL.

### The nAngptl4^E40K^ variant reshapes residue 40 engagement with LPL and oligomerization interfaces

To test our hypothesis that nAngptl4^E40K^ binds more weakly than nAngptl4^WT^ to LPL, we performed simulations of nAngptl4^E40K^ in complex with LPL for direct comparison with our earlier nAngptl4^WT^–LPL simulations. As in those simulations, a representative nAngptl4^E40K^ structure was docked to an open-lid LPL conformation, and the top-scoring biologically relevant model—selected based on known binding sites^47^—was simulated in triplicate (1 µs each). Overall, nAngptl4^E40K^ behaved similarly to nAngptl4^WT^ in complex with LPL. RMSD profiles showed only minor fluctuations, indicating stable complexes (**Supplementary Fig. 12A**), and both variants exhibited only slight reorientations across the top of LPL relative to the initial pose (**Supplementary Fig. 12B–C**). These reorientations were consistent between nAngptl4^WT^ and nAngptl4^E40K^, with similar residues on both proteins frequently engaging LPL (**Supplementary Fig. 12D, F–G**) and comparable overall binding interaction energies (**Supplementary Fig. 12E**).

Subtle differences emerged in the variants’ functional interactions with LPL. SASA measurements revealed that nAngptl4^WT^ provided greater lid coverage than nAngptl4^E40K^ **(Fig. 5A)**, while both exhibited minimal lipid pore coverage (**Fig. 5B**). These differences corresponded with modest variations in lid dynamics, as some E40K simulations showed earlier and larger fluctuations in lid closure. While their total binding energetics were similar, decomposition analysis showed that nAngptl4^WT^ engaged approximately twice as many LPL residues when bound as nAngptl4^E40K^ (**Fig. 5D**; **Supplementary Fig. 13A–C**). Notably, residue E40 contributed to LPL binding in nAngptl4^WT^, whereas K40 did not in the mutant, mirroring trends observed for H1 versus H1-E40K. This difference is consistent with prior observations that E40 participates in more transient salt bridge interactions, both within nAngptl4^WT^ and with LPL, whereas K40 appears largely restricted to interactions with residue 39 in nAngptl4^E40K^. Together, these results suggest that reduced energetic contributions in E40K may weaken its engagement with LPL.

**Figure 5.**
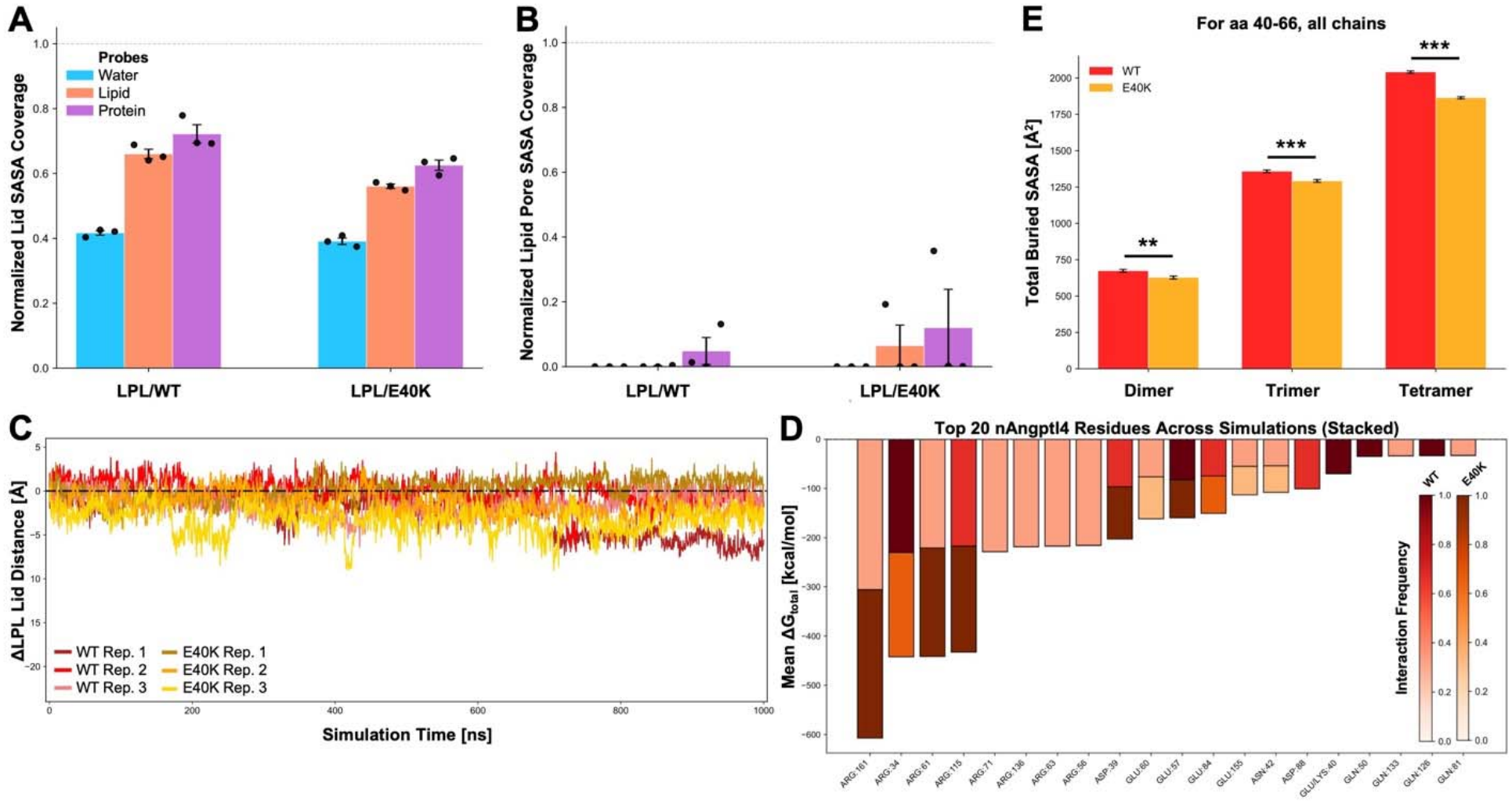
nAngptl4^WT^ restrains LPL lid dynamics more effectively than loss-of-function variant nAngptl4^E40K^. (A–B) Solvent accessible surface area (SASA) coverage of LPL lid residues (A) and lipid pore entrance-surrounding residues (B) by simulated nAngptl4 variants from 900–1000 ns, normalized to unbound LPL with values ranging from 0 to 1, where 0 indicates no coverage and 1 indicates complete coverage. Probe radii correspond to water, lipid, and protein (0.14, 0.50, and 1.00 nm, respectively). Mean values (black bars) are shown with standard error of the mean (SEM). (C) Change in LPL lid distance over simulation time relative to the initial distance. The dashed line indicates Δ = 0. (D) Top 20 energetic contributors among nAngptl4 residues to LPL binding. ΔG_bind_ is shown per variant, with interaction frequency indicated by color. (E) Total buried SASA of residues 40–66 across all chains in nAngptl4 oligomers derived from docking. Error bars represent standard deviation. Statistical significance across oligomeric pairs in nAngptl4 variants was assessed using Welch’s t-test (** = P<0.01, *** = P<0.001; n = 500).

Because residue 40 lies adjacent to the primary LPL-binding site, we next considered how this region might be influenced by nAngptl4 oligomerization. Beyond monomer stability, impaired oligomerization^44^ has been proposed as a mechanism underlying reduced LPL inhibition by nAngptl4^E40K^. Although the structural determinants of higher-order nAngptl4 assembly remain unclear, early studies implicated disulfide bonds between C76 and C80 in the hinge region^41,44^, while later work suggested oligomerization can persist even if these bonds are disrupted^25,43^. In the absence of a defined dimer interface, we performed preliminary blind docking using representative monomer structures from the late stages of the simulations for each variant (**Fig. 4B**). The top 50 scoring models were analyzed to quantify predicted monomer–monomer interfacial contacts (**Supplementary Fig. 14A–C**).

Putative nAngptl4^WT^ dimers showed greater involvement of hinge and terminal regions at the interface, whereas nAngptl4^E40K^ dimers more frequently engaged central helical regions. Consistent with this, SASA analysis across docked oligomer models revealed that the primary LPL-binding site was more frequently engaged in nAngptl4^WT^ dimers, trimers, and tetramers (**Fig. 5E**). This suggests that, in addition to mediating LPL binding, this region may contribute to oligomer formation and stabilization. While limited by the use of rigid docking and static conformations, these results indicate that a single point mutation can shift preferred modes of self-association, potentially altering the balance between oligomer stability and LPL binding. Together, these findings suggest that E40K does not grossly destabilize the monomer but instead subtly reshapes nAngptl4’s interaction networks and self-association interfaces.

## Discussion

Here, we present the first molecular dynamics (MD) investigation of nAngptl4, using peptide fragments and analysis of clinically relevant loss-of-function variants to uncover novel mechanistic insights into nAngptl4-mediated LPL inhibition. While nAngptl4 has long been known to suppress LPL activity^31,41,42^, the structural basis of this inhibition has remained unclear. MD simulations have allowed us to capture the complex structural conformations of LPL and how these dynamics affect its regulation by specific cofactors. Our results support a model in which nAngptl4 inhibits LPL primarily by binding to the lid and restricting its dynamics, rather than directly occluding the lipid pore. This suggests that early-stage inhibition is driven by modulation of conformational flexibility, with destabilization occurring over longer timescales.

This mechanism refines prior models that emphasize LPL destabilization^23,25^ and dimer dissociation^25,27–30^. Consistent with HDX-MS data^46,47^, our simulations reveal that nAngptl4 binds across LPL’s lid and adjacent regions spanning the N-terminal domain, positioning it to influence both catalytic accessibility and structural integrity. Notably, solvent accessibility of the lipid pore remains largely unchanged upon nAngptl4 binding, while lid accessibility and dynamics are significantly reduced, supporting a model in which constrained lid motion impairs substrate processing and LPL’s structure–function relationships. Furthermore, these ideas of initially regulating LPL’s activity via its lid dynamics and prevention of LPL C-terminal domain flipping—a behavior observed in unbound LPL in our previous work^61^—remain consistent with our proposed biphasic model of initially reversible inhibition^45^ by nAngptl4, followed by irreversible, global destabilization of LPL^23,25^.

Our *in silico* peptide-based analyses further demonstrate that binding mode dictates the inhibitory mechanism. Full-length nAngptl4 utilizes its coiled-coil domain to preferentially engage the lid of LPL, whereas nAngptl4-derived peptides bind more variably across LPL’s key functional domains but also cluster around the lid. Notably, these peptides represent a mechanistically distinct alternative function of nAngptl4, despite containing the exact same residues. All three single-coil peptides significantly activated LPL above its baseline-level, and reduced nAngptl4-mediated inhibition. This striking observation highlights remaining questions surrounding LPL regulation. For example, what structural and biochemical features are critical for mediating LPL activity across a spectrum from inhibition to activation? Moreover, as we begin to explore therapeutic routes for modulating LPL, this motivates the consideration of endogenous inhibitors and loss-of-function variants, rather than limiting design to known activators. The consistency of the peptides, especially H1-E40K, to increase LPL hydrolysis, retain LPL lid flexibility, and reduce the effect of nAngptl4 serves as a useful starting point in exploring this potential therapeutic design strategy.

Our simulations also provide insight into the molecular basis of the E40K loss-of-function variant of nAngptl4. Rather than causing global destabilization, we find that the E40K mutation induces subtle changes in local interaction networks centered on residue 40. In the wild-type protein, E40 participates in transient salt-bridge interactions both within nAngptl4 and with LPL, and exhibits greater flexibility across the secondary binding site within full-length Helix 2. In contrast, K40 forms more restricted interactions, both structurally and electrostatically. These differences extend to LPL binding, where fewer residues contribute to binding in the E40K variant, despite a similar overall binding affinity for LPL as the wild-type construct, suggesting a redistribution of interfacial contacts rather than a complete loss of binding upon mutation. Preliminary analyses further suggest that E40K alters preferred nAngptl4 self-association interfaces, consistent with a biphasic model in which the mutation weakens early-stage LPL engagement and may impair inhibition through altered oligomerization over longer timescales.

Several limitations should be considered when interpreting these results. MD simulations are inherently constrained by accessible timescales, and processes such as nAngptl4 oligomer destabilization—particularly for the E40K variant—or nAngptl4-mediated destabilization of LPL likely occur beyond the microsecond scale sampled here. In addition, our simulations do not fully capture the biological context of LPL, which is typically membrane-associated and interacts with receptors^15–17,87,88^, proteoglycans^89,90^, lipoproteins^16^, and extracellular lipids^91^. Additional structural features, including N-linked glycans^92,93^, Ca^2+^ coordination at the active site^94^, and potential proteolytic processing between LPL domains^95^, were also not explicitly modeled. Furthermore, the use of rigid protein–protein docking limits exploration of conformational variability and may bias initial binding poses. However, subsequent MD simulations of these docked complexes allowed for relaxation and reorientation of these structures. Both peptides and full-length nAngptl4 sampled alternative binding configurations or dissociated entirely within the simulated timescales, suggesting that the observed interactions are physically reasonable.

We also acknowledge that simulating LPL and nAngptl4 as monomers may not fully capture their most biologically relevant forms, as both proteins are known to adopt functionally active oligomeric states^16,18–21,23,25,41,43–45^. However, prior studies have demonstrated that each can exist and remain active as monomers^15,16,23,44,45^. Moreover, the precise interfaces governing self-association—particularly in the context of LPL–nAngptl4 interactions—remain poorly defined, making accurate simulation of higher-order complexes challenging. This limitation is especially pronounced for nAngptl4, for which no experimental structure is currently available. Therefore, we elected to model a simplified 1:1 complex to isolate key interaction features and obtain atomistic insights without introducing artifacts arising from potentially incorrect oligomerization. Despite these limitations, our findings provide atomistic insights into LPL inhibition that are difficult to obtain experimentally. As computational resources continue to expand, incorporating higher-order oligomers, more complete biological environments, and longer simulation timescales will further refine our understanding of LPL regulation.

Moreover, the use of peptide fragments derived from endogenous proteins represents a promising avenue for therapeutic targeting of LPL. These findings expand upon our previous work^82^, which demonstrated that an ApoC-II-derived peptide interacts with the LPL lid domain to stabilize the enzyme while increasing lid flexibility and catalytic activity. Here, we tested for the first time whether peptides encompassing the LPL-binding region of nAngptl4 could recapitulate the inhibitory properties of the full-length protein. Surprisingly, all peptides exhibited dual activity, attenuating nAngptl4-mediated inhibition while directly enhancing LPL activity. Although the underlying mechanisms require further study, these effects may reflect the relatively unstructured nature of the peptides compared with the predominantly helical full-length protein, together with the complex network of LPL interactions surrounding the lid domain. The ability of these peptides to block nAngptl4-mediated inhibition suggests that their interaction with LPL prevents engagement of the full-length nAngptl4 with the lid domain. Defining the relative contributions of peptide structure, sequence composition, and stability will be critical for guiding the development of peptide-based LPL therapeutics.

This work further addresses a longstanding question in LPL biology: how LPL is regulated across its monomeric and dimeric states, and whether these forms differ mechanistically in their function. Experimental studies have yet to resolve whether activity differs substantially between monomers and dimers, an issue that is particularly relevant given the multiple modes by which nAngptl4 may inhibit LPL. One possibility is that LPL is more stable in its dimeric form, while nAngptl4 promotes dissociation into monomers that, although still catalytically active, are more susceptible to irreversible destabilization. Binding of nAngptl4 to monomeric LPL may then facilitate unfolding of the hydrolase domain, particularly in the absence of stabilizing interactions with receptors such as GPIHBP1. At the same time, the diverse set of LPL regulators, including receptors, activators, and inhibitors, across tissues suggests that these mechanisms are likely context-dependent and may vary with physiological or disease state.

Overall, this work highlights the importance of LPL lid dynamics and distributed interaction networks in LPL inhibition. By performing the first physics-based simulations of nAngptl4 in complex with LPL, we have identified novel mechanisms of inhibition and elucidated the nuanced impact of a well-characterized loss-of-function variant^49^. These results provide a framework for understanding nAngptl4-mediated regulation of LPL and may inform future therapeutic strategies targeting lipid metabolism.

## Acknowledgments

This work utilized the Alpine high-performance computing resource and Blanca condo computing resource at the University of Colorado Boulder. Alpine is funded through a joint effort by the University of Colorado Boulder, the University of Colorado Anschutz, and Colorado State University. Blanca receives funding from its computing users and the University of Colorado Boulder. This work also utilized resources at the Structural Biology and Biophysics Core Facilities at the University of Colorado Anschutz School of Medicine, which is supported in part by the School of Medicine.

