## Supplemental Information for "Lid Restriction Underlies Early nAngptl4-Mediated Lipoprotein Lipase Inhibition and Is Prevented by nAngptl4^E40K^ and Peptide Mimetics"

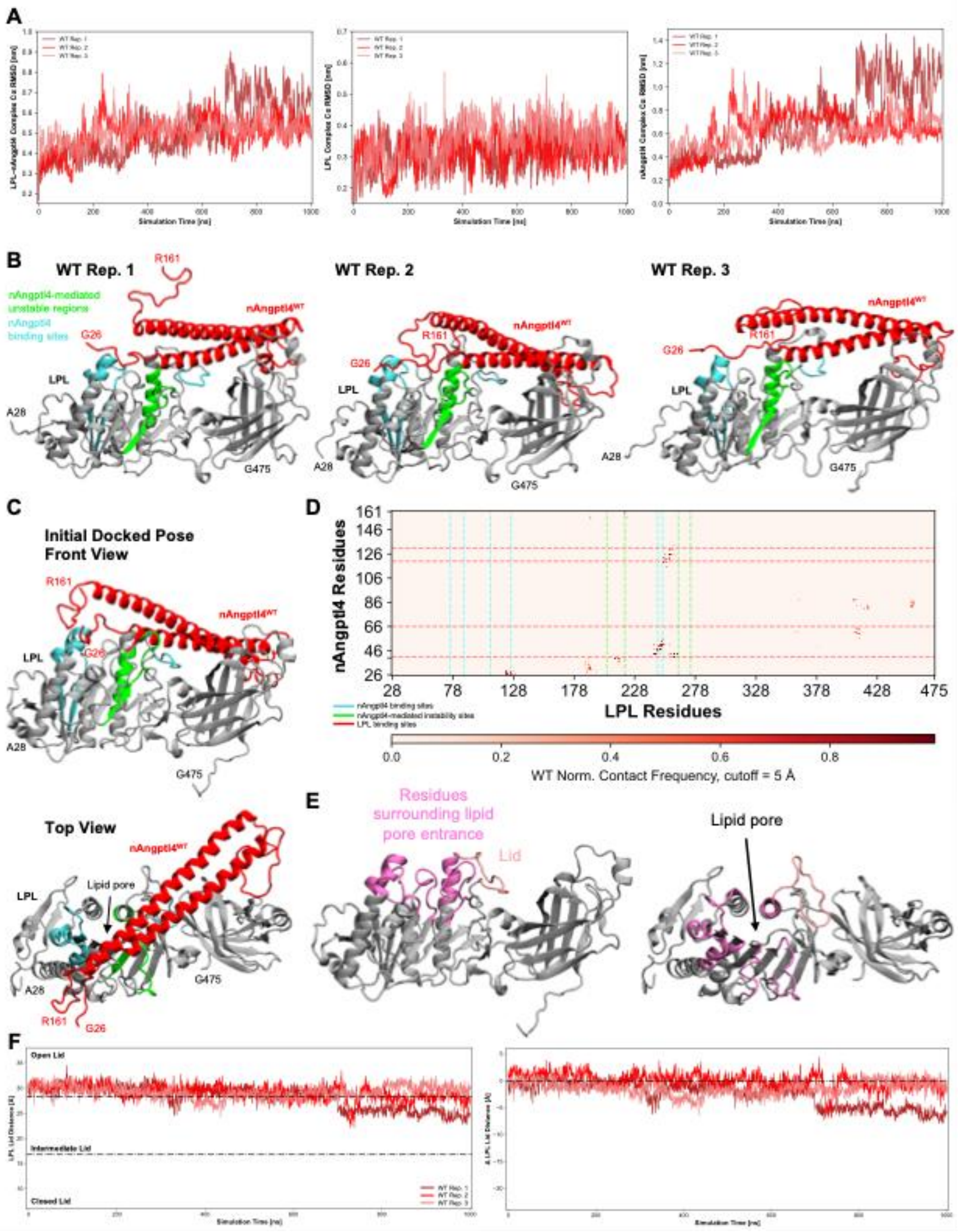


**Supplementary Figure 1.** (A) RMSD of Cα backbone of the LPL–nAngptl4$^{WT}$ complex (left), LPL within the complex (center), and nAngptl4$^{WT}$ within the complex (right) over simulation time. (B) Front view of LPL bound to nAngptl4$^{WT}$ (final simulation time point, t = 1000 ns) for the three simulations. LPL is seen in gray, with the nAngptl4 binding sites in cyan (aa 76-87, 109-126, 245-251; LPL numbered 1-475) and allosterically unstable regions in green (aa 205-220, 264-274). nAngptl4$^{WT}$ is seen in red. (C) Initial docked pose for simulations with the front view (top) and top view (bottom). Protein colors are the same as described in (B). (D) Normalized (Norm.) contact map of nAngptl4$^{WT}$ (WT) and LPL residues across the three simulations, from 900-1000 ns. Dashed lines represent bounds for binding sites on LPL and nAngptl4, for one another. (E) Front (left) and top (right) views of LPL with the residues surrounding the lipid pore entrance (medium pink) and lid (light pink) shown. Pore-surrounding residues include: aa 81-91, 114-128, 159-161, 183-192, 210-223, 261-269; lid residues include: aa 245-260. (F) Left: LPL Lid distance over simulation time. Lid distance clusters are taken from previous work. Black dash-dot lines represent cluster bounds. Right: Change in LPL lid distance over simulation time, compared to its initial distance. Black dash-dot line represents Δ = 0.

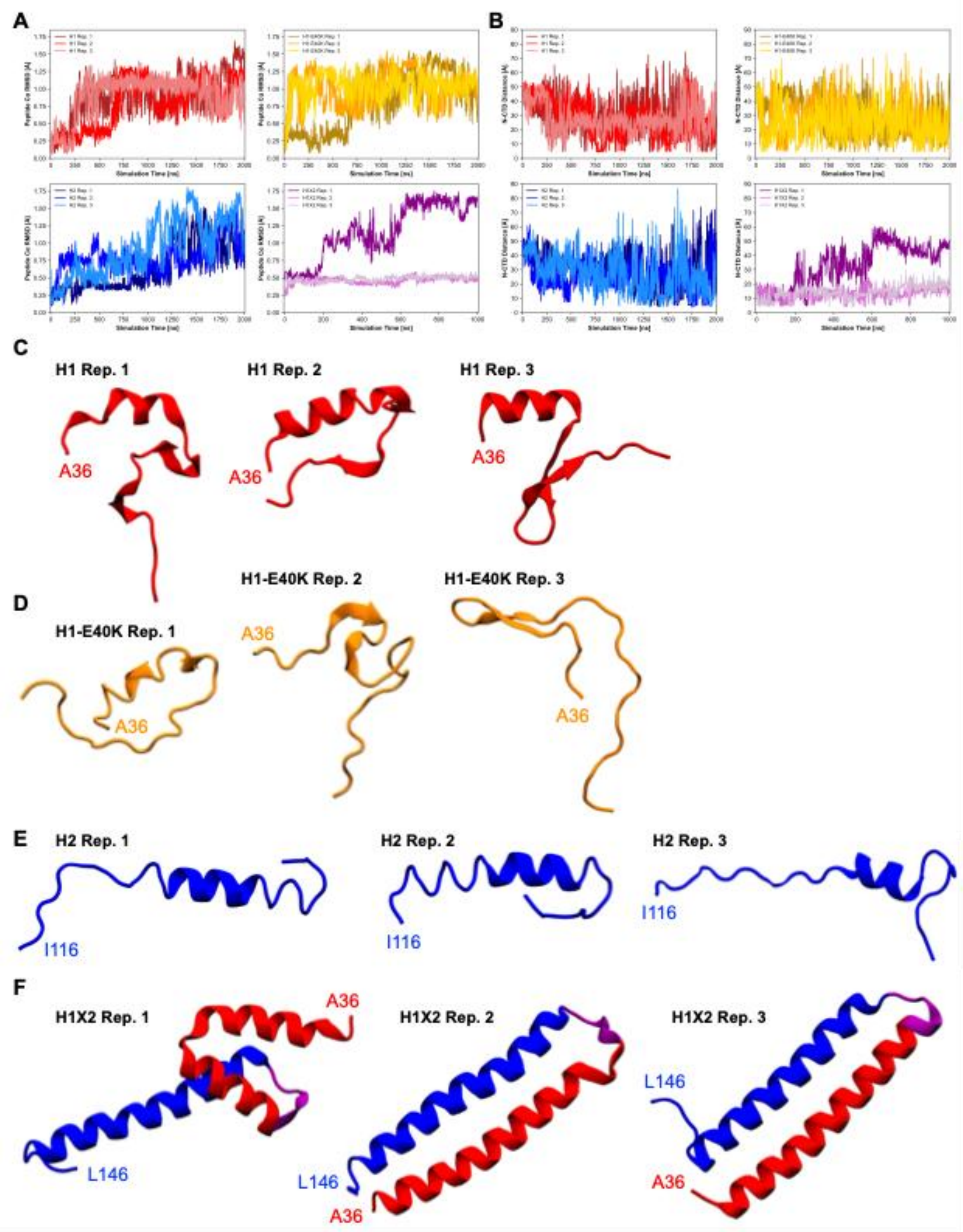


**Supplementary Figure 2**. (A) RMSD of Cα backbone of Angptl4 peptides over simulation time. H1 is shown in reds, H1-E40K in oranges, H2 in blues, and H1X2 in purples. (B) NTD-CTD terminal residue distances of Angptl4 peptides over simulation time. H1 is shown in reds, H1-E40K in oranges, H2 in blues, and H1X2 in purples. (C) Final simulation trajectory snapshots (t = 2000 ns) for the H1 replicates. (D) Final simulation trajectory snapshots (t = 2000 ns) for the H1-E40K replicates. (E) Final simulation trajectory snapshots (t = 2000 ns) for the H2 replicates. (F) Final simulation trajectory snapshots (t = 1000 ns) for the H1X2 replicates. H1 residues are seen in red, the 5-residue linker in purple, and H2 residues in blue.

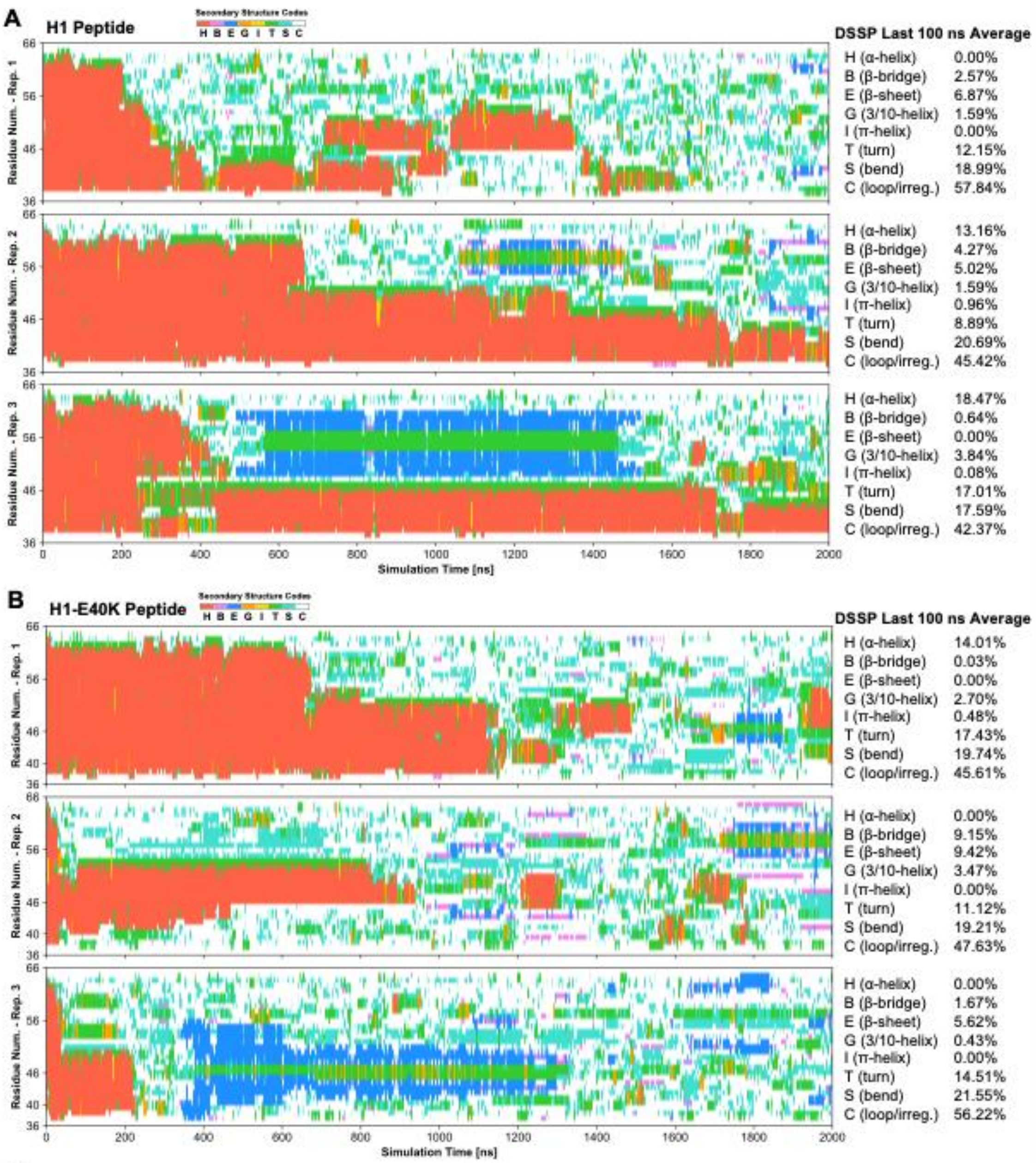


**C**

| H1 Average (DSSP) | | BeStSel Estimation | |
|---|---|---|---|
| H (α-helix) | 10.54% | Helix | 10.54% |
| B (β-bridge) | 2.49% | Anti/Parallel | 3.96% |
| E (β-sheet) | 3.96% | Turn | 12.68% |
| G (3/10-helix) | 2.43% | Others | 72.90% |
| I (π-helix) | 0.35% | | |
| T (turn) | 12.68% | | |
| S (bend) | 19.09% | | |
| C (loop/irreg.) | 48.54% | | |

| H1-E40K Average (DSSP) | | BeStSel Estimation | |
|---|---|---|---|
| H (α-helix) | 4.67% | Helix | 4.67% |
| B (β-bridge) | 3.62% | Anti/Parallel | 5.01% |
| E (β-sheet) | 5.01% | Turn | 14.35% |
| G (3/10-helix) | 2.20% | Others | 75.97% |
| I (π-helix) | 0.16% | | |
| T (turn) | 14.35% | | |
| S (bend) | 20.17% | | |
| C (loop/irreg.) | 49.82% | | |

**Supplementary Figure 3.** (A) DSSP profiles for H1 Peptide simulations over simulation time. Secondary structure codes include: α-helix (H), residue in isolated β-bridge (B), extended strand participating in β-ladder/sheet (E), 3/10-helix (G), π-helix (I), hydrogen-bonded turn (T), bend (S), and loop/irregular elements (C). DSSP values averaged over the last 100 ns/simulation are shown. (B) DSSP profiles for H1-E40K Peptide simulations over simulation time. Secondary structure codes are the same as in (A). DSSP values averaged over the last 100 ns/simulation are shown. (C) The replicate average for H1 (left) and H1-E40K (right) are shown, with the values converted to a BeStSel estimation.

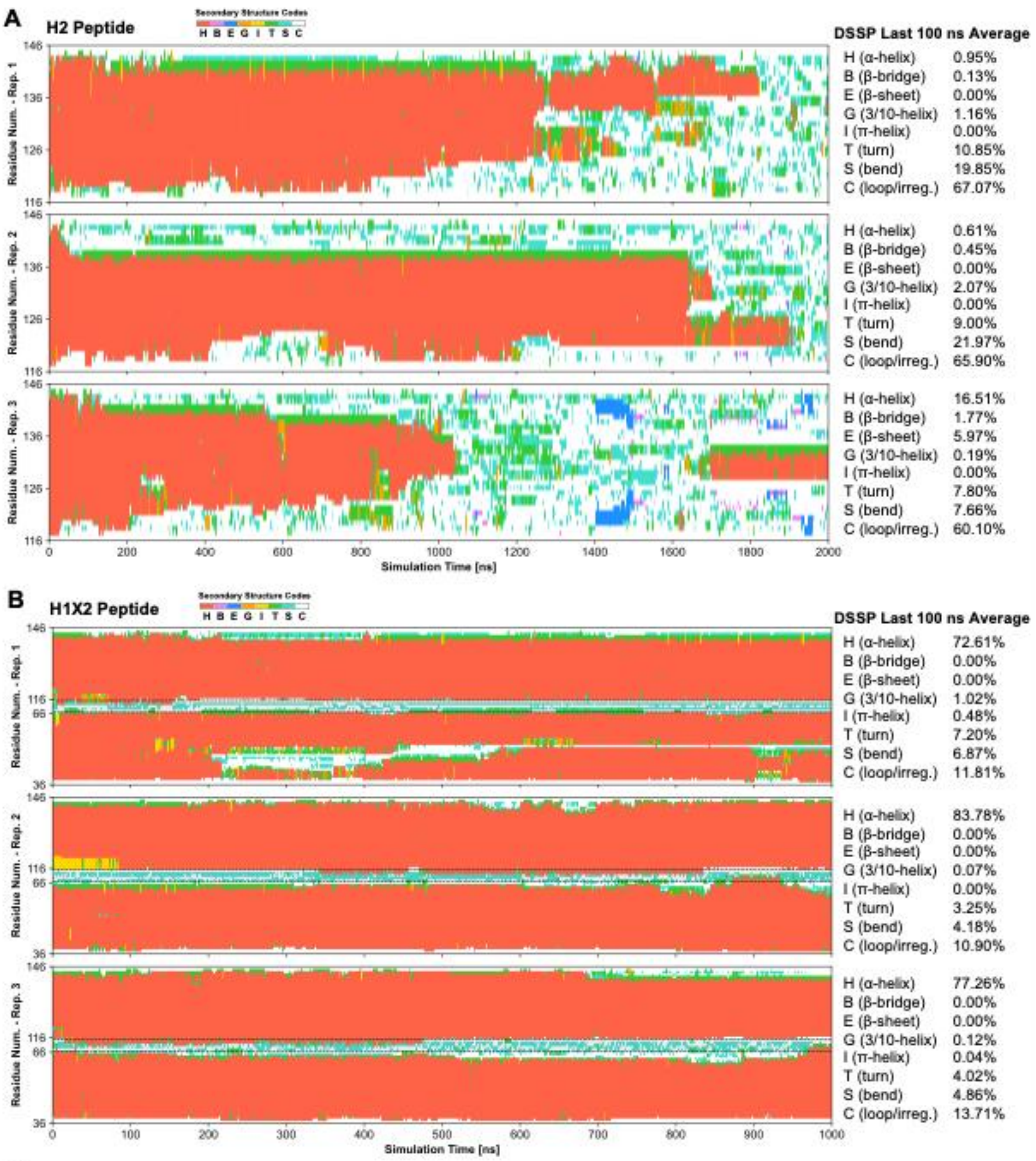


C

| H2 Average (DSSP) | |
|---|---|
| H (α-helix) | 6.02% |
| B (β-bridge) | 0.78% |
| E (β-sheet) | 1.99% |
| G (3/10-helix) | 1.14% |
| I (π-helix) | 0.00% |
| T (turn) | 9.22% |
| S (bend) | 16.49% |
| C (loop/irreg.) | 64.36% |

| BeStSel Estimation | |
|---|---|
| Helix | 6.02% |
| Anti/Parallel | 1.99% |
| Turn | 9.22% |
| Others | 82.77% |

| H1X2 Average (DSSP) | |
|---|---|
| H (α-helix) | 77.88% |
| B (β-bridge) | 0.00% |
| E (β-sheet) | 0.00% |
| G (3/10-helix) | 0.40% |
| I (π-helix) | 0.17% |
| T (turn) | 4.82% |
| S (bend) | 5.30% |
| C (loop/irreg.) | 12.14% |

**Supplementary Figure 4.** (A) DSSP profiles for H2 Peptide simulations over simulation time. Secondary structure codes include: α-helix (H), residue in isolated β-bridge (B), extended strand participating in β-ladder/sheet (E), 3/10-helix (G), π-helix (I), hydrogen-bonded turn (T), bend (S), and loop/irregular elements (C). DSSP values averaged over the last 100 ns/simulation are shown. (B) DSSP profiles for H1X2 Peptide simulations over simulation time. Secondary structure codes are the same as in (A). DSSP values averaged over the last 100 ns/simulation are shown. (C) The replicate average for H2 (left) and H1X2 (right) are shown, with the values converted to a BeStSel estimation for H2.

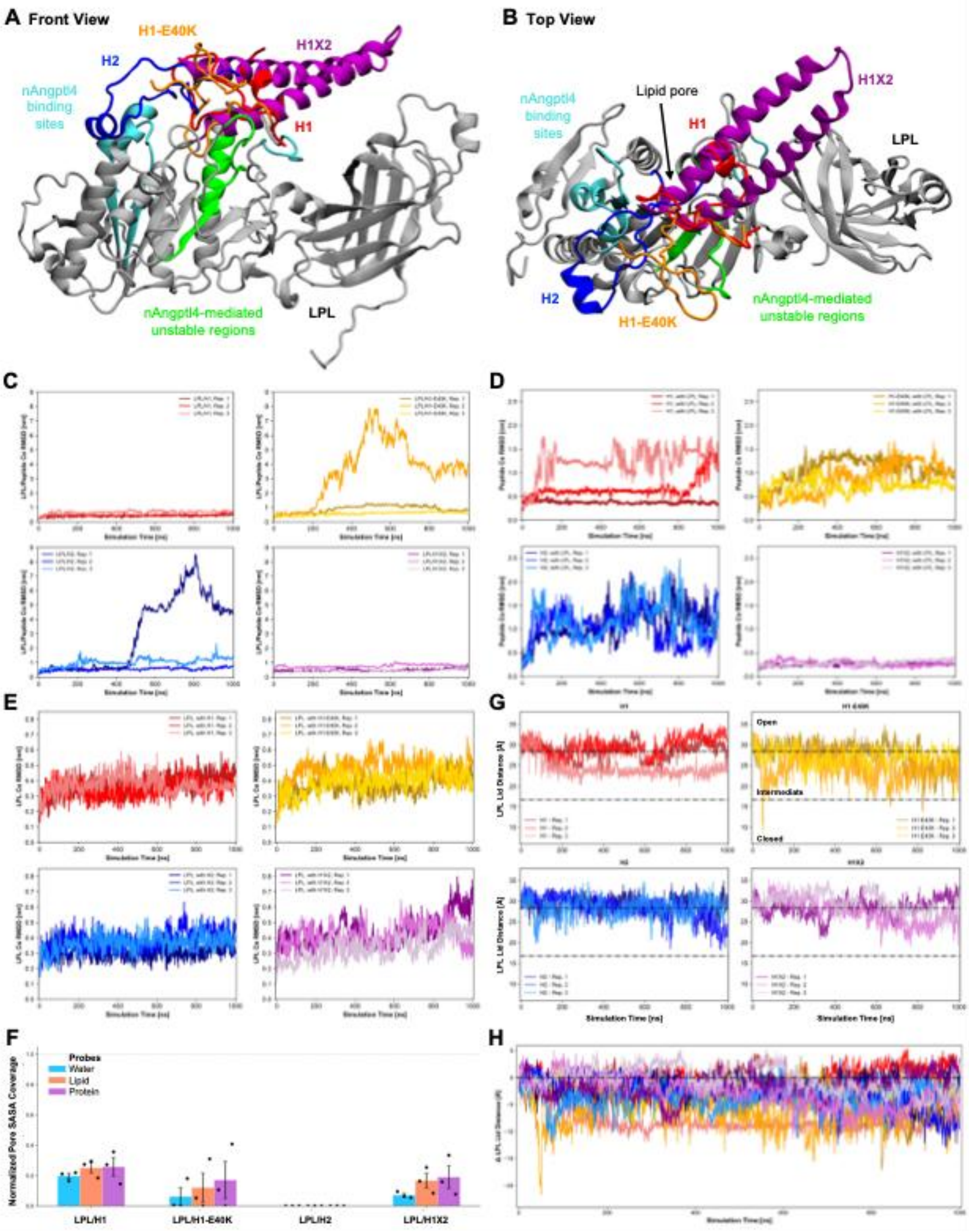


**Supplementary Figure 5.** (A) Front view of open-lid LPL docked to Angptl4 peptides. LPL is seen in gray, with the nAngptl4 binding sites in cyan (aa 76-87, 109-126, 245-251; LPL numbered 1-475) and allosterically unstable regions in green (aa 205-220, 264-274). H1, H1-E40K, H2, and H1X2 are seen in red, orange, blue, and purple, respectively. (B) Top view of peptides docked to LPL, with the same color descriptions as in (A). (C) RMSD of Cα backbone of the LPL + Peptide complexes over simulation time. H1 is shown in reds, H1-E40K in oranges, H2 in blues, and H1X2 in purples. (D) RMSD of Cα backbone of Angptl4 peptides over simulation time. H1 is shown in reds, H1-E40K in oranges, H2 in blues, and H1X2 in purples. (E) RMSD of Cα backbone of LPL over simulation time. H1 is shown in reds, H1-E40K in oranges, H2 in blues, and H1X2 in purples. (F) Normalized solvent accessible surface area (SASA) coverage of peptides on LPL for lid residues from 900-1000 ns. Coverage was normalized against ligand-free LPL, ranging from [0,1], with 0 representing no coverage on an LPL region and 1 representing complete coverage (with respect to time and residues). Probes represent water, lipid, and protein sizes (radii = 0.14, 0.50, and 1.00 nm, respectively). Mean coverage value is shown with standard error of the mean (SEM) in black bars. G) LPL Lid distance over simulation time. Lid distance clusters are taken from previous work. Black dash-dot lines represent cluster bounds. Bounds are listed on the top, right plot (H1-E40K). (H) Change in LPL lid distance over simulation time, compared to its initial distance. Black dash-dot line represents $\Delta = 0$.

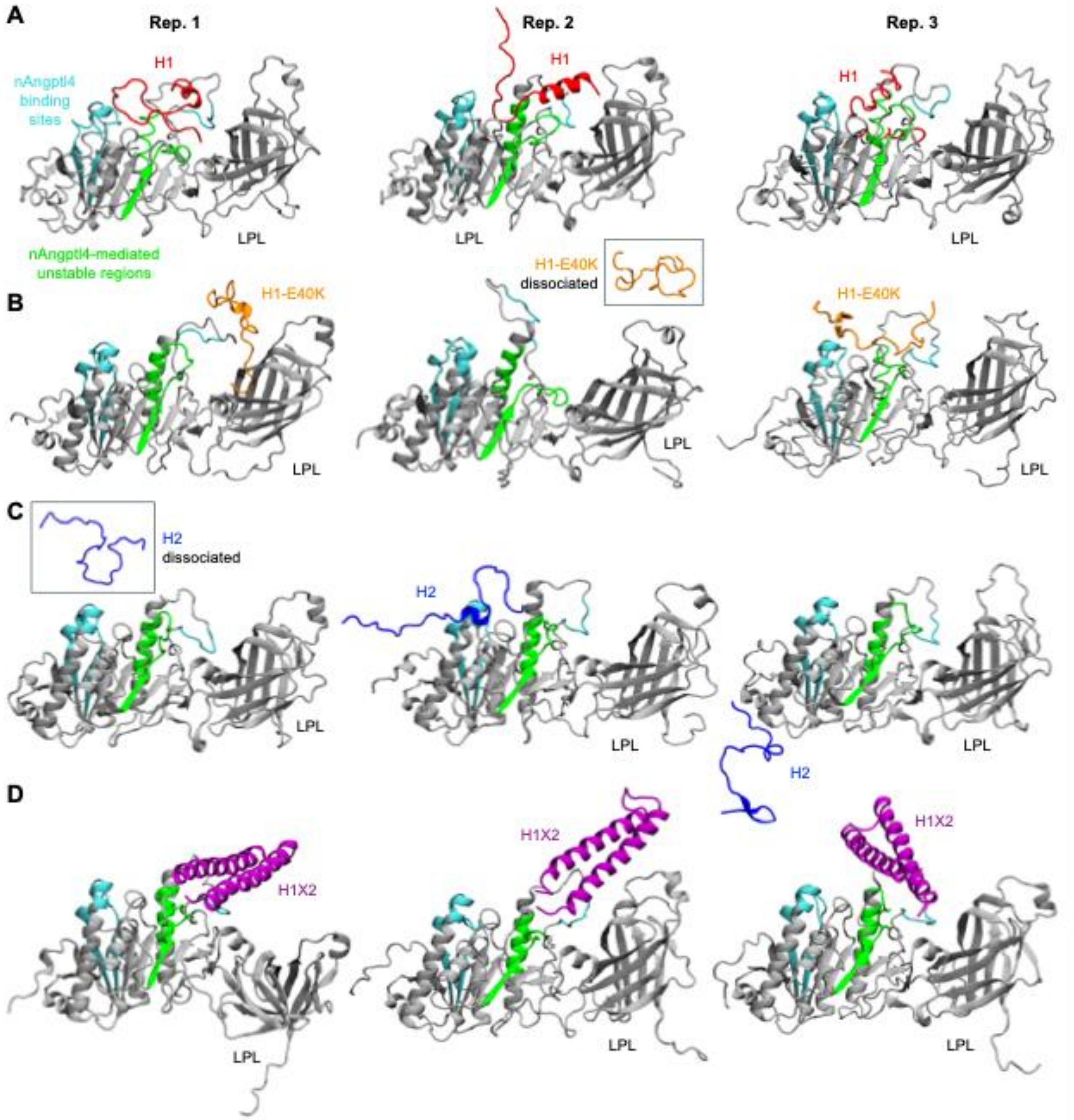


**Supplementary Figure 6.** (A) Front view of LPL–H1 simulations, as the final simulation trajectory snapshots (t=1000 ns) for the replicates. LPL is seen in gray, with the nAngptl4 binding sites in cyan (aa 76-87, 109-126, 245-251; LPL numbered 1-475) and allosterically unstable regions in green (aa 205-220, 264-274). H1 is seen in red. Snapshots for Reps. 1, 2, and 3 are seen from left, middle, to right, resp. (B) Front view of LPL–H1-E40K simulations, as the final simulation trajectory snapshots (t=1000 ns) for the replicates. LPL colors are described in (A). H1-E40K is seen in orange. In Rep. 2, H1-E40K dissociated, so final structures of the separated LPL and H1-E40K are shown. (C) Front view of LPL–H2 simulations, as the final simulation trajectory snapshots (t=1000 ns) for the replicates. LPL colors are described in (A). H2 is seen in blue. In Rep. 1, H2 dissociated, so final structures of the separated LPL and H2 are shown. (D) Front view of LPL–H1X2 simulations, as the final simulation trajectory snapshots (t=1000 ns) for the replicates. LPL colors are described in (A). H1X2 is seen in purple. In Rep. 1, the transition of stable LPL to an unstable form, via CTD flipping, is observed.

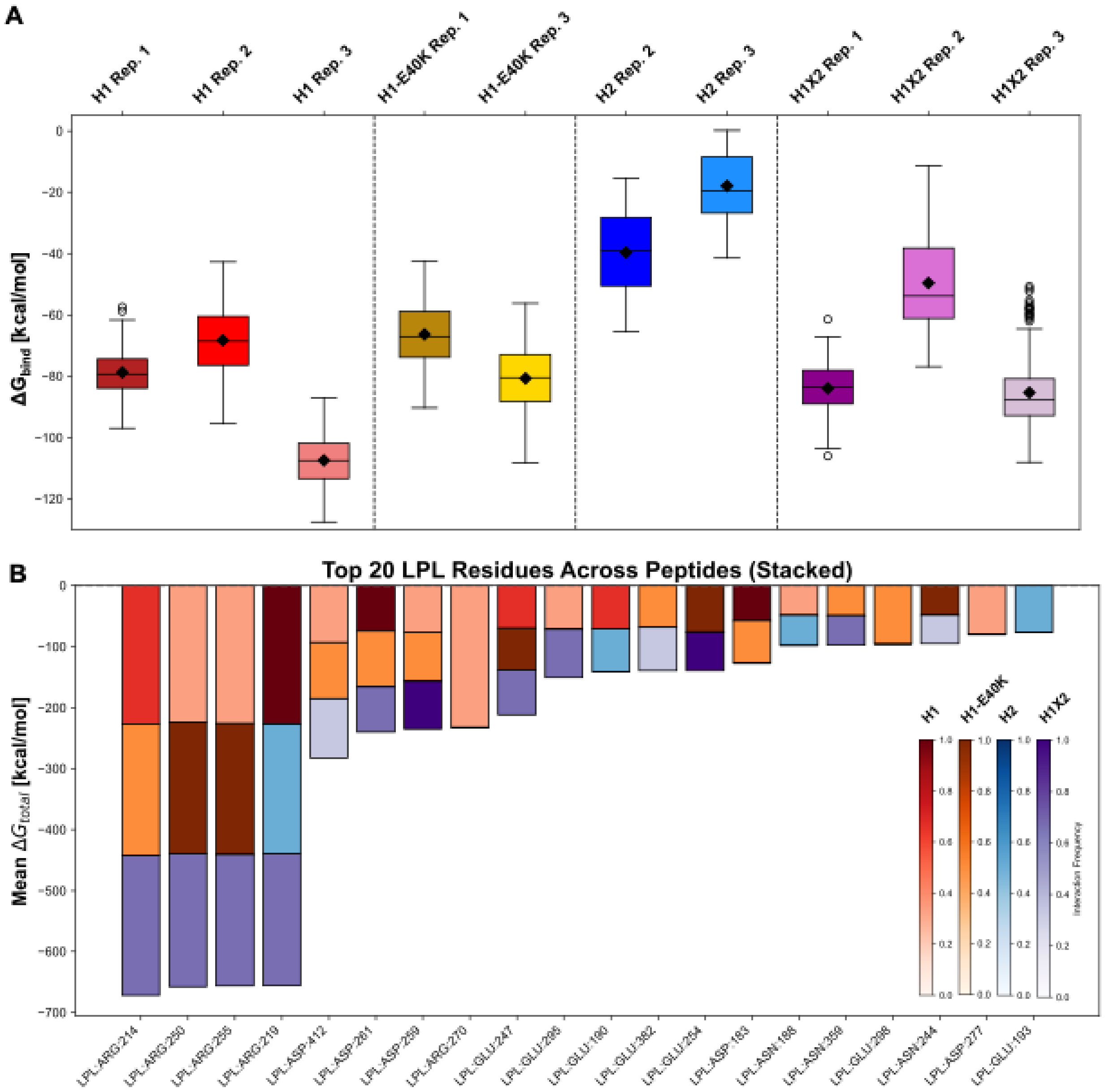


**Supplementary Figure 7.** (A) Binding interaction energies ($\Delta G_{bind}$) between peptides and LPL from 900-1000 ns per simulation. The standard deviation, mean (diamond) and median (line) are shown. H1-E40K Rep. 2 and H2 Rep. 1 are not shown since the peptides dissociated from LPL. (B) Summaries of binding energetic contributions across the top 20 LPL residues in peptide interactions. The $\Delta G_{bind}$ value per peptide is shown, with the interaction frequency as the color. Interaction frequency scale bars are shown on the right, per peptide.

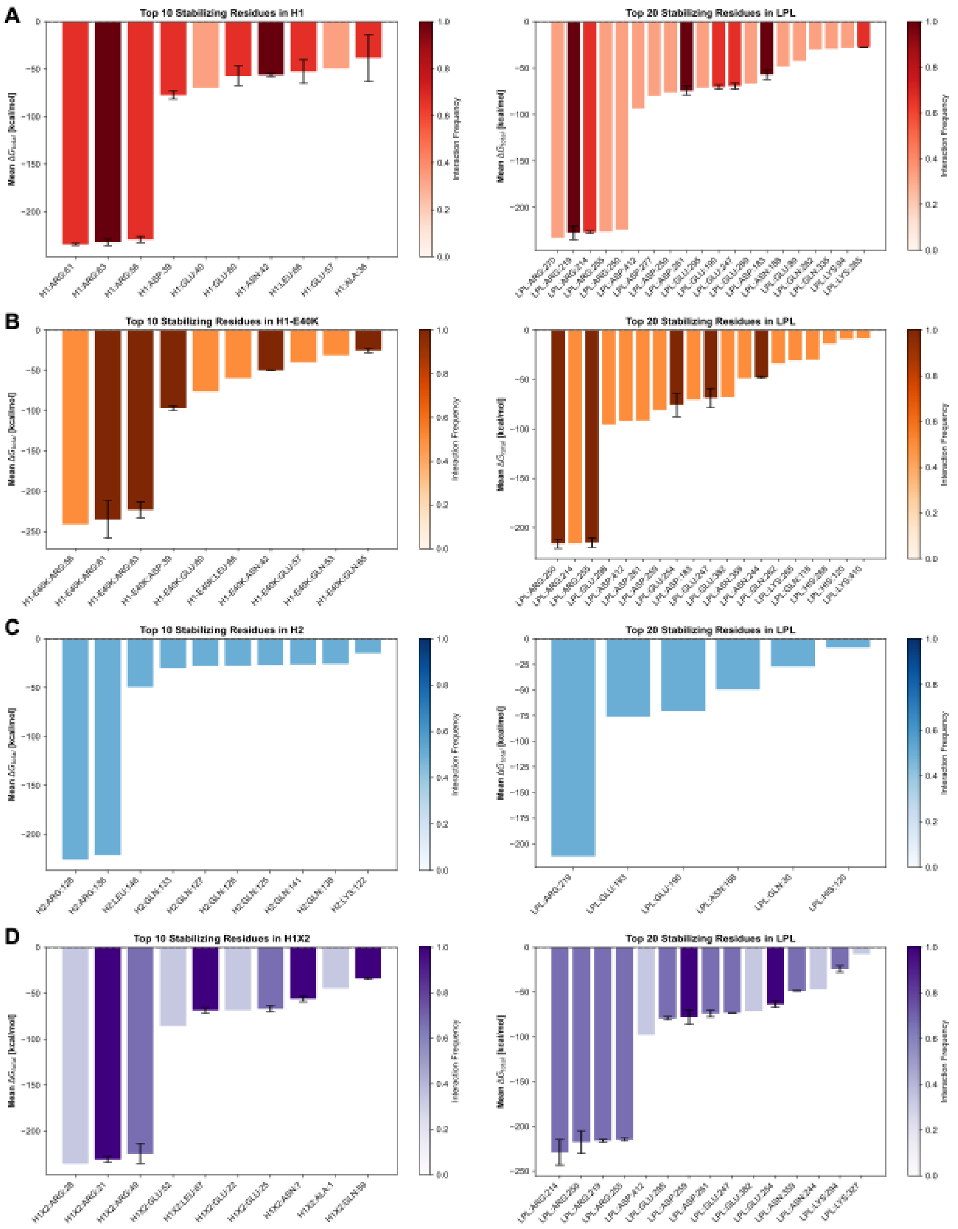


**Supplementary Figure 8.** (A) Summaries of binding energetic contributions across the top 10 H1 residues (left) and top 20 LPL residues (right). The mean $\Delta G_{bind}$ value across the replicates is shown with the standard deviation for residues involved in all 3 simulations. Colors correspond to the interaction frequency across the simulations, with scale bars shown on the right. (B) Summaries of binding energetic contributions across the top 10 H1-E40K residues (left) and top 20 LPL residues (right). The mean $\Delta G_{bind}$ value across the replicates is shown with the standard deviation for residues involved in all 3 simulations. Colors correspond to the interaction frequency across the simulations, with scale bars shown on the right. (C) Summaries of binding energetic contributions across the top 10 H2 residues (left) and top 6 LPL residues, as only 6 were determined (right). The mean $\Delta G_{bind}$ value across the replicates is shown with the standard deviation for residues involved in all 3 simulations. Colors correspond to the interaction frequency across the simulations, with scale bars shown on the right. (D) Summaries of binding energetic contributions across the top 10 H1X2 residues (left) and top 20 LPL residues (right). The mean $\Delta G_{bind}$ value across the replicates is shown with the standard deviation for residues involved in all 3 simulations. Colors correspond to the interaction frequency across the simulations, with scale bars shown on the right. Residue numbers in H1X2 are shifted, and correspond to the following in the original NAngptl4 sequence, from left to right: R63, R56, R128, E131, L146, E57, E60, N42, A36, Q138.

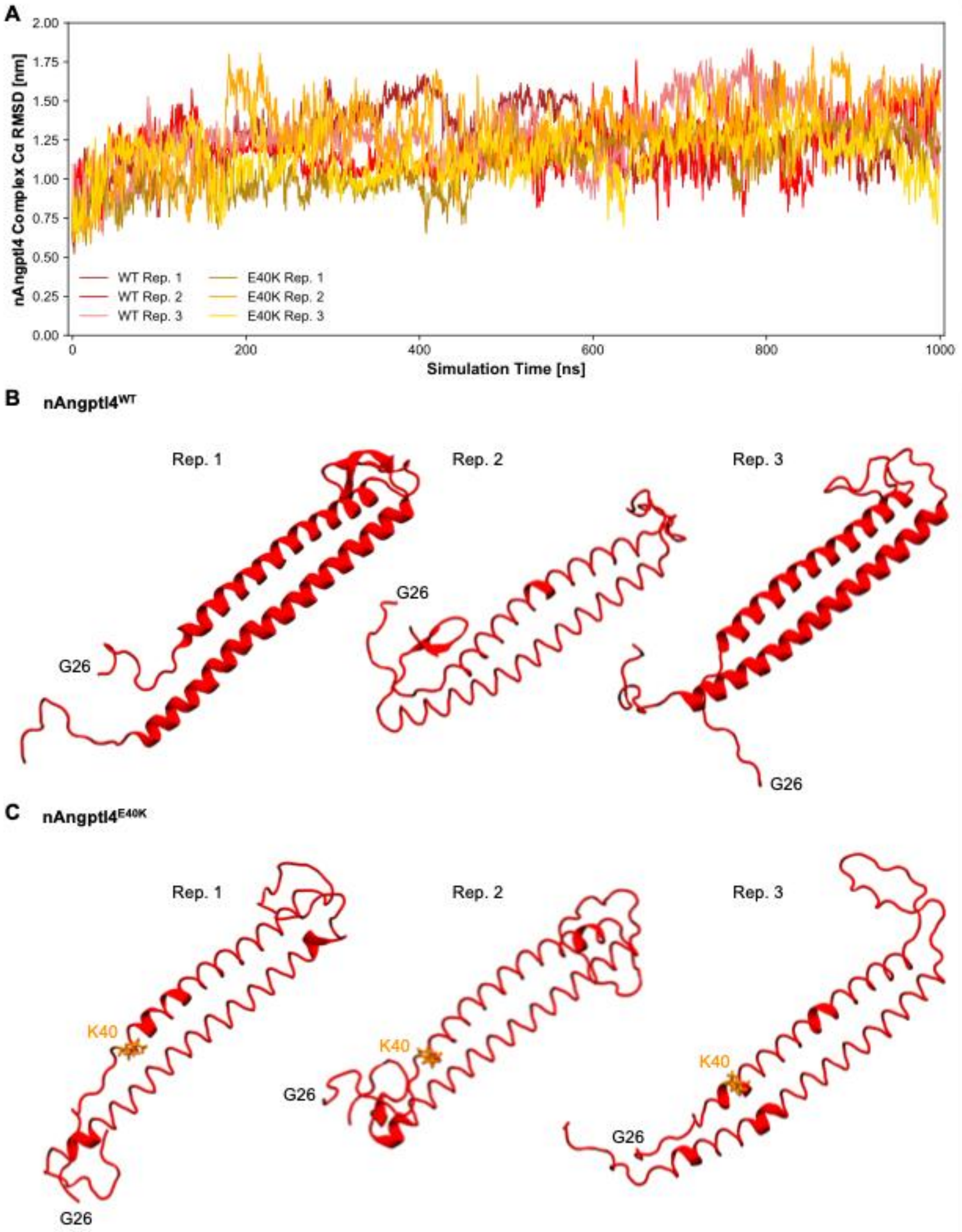


**Supplementary Figure 9.** (A) RMSD of Cα backbone of nAngptl4$^{WT}$ (reds) and nAngptl4$^{E40K}$ (oranges) over simulation time. (B) Final simulation trajectory snapshots (t = 1000 ns) for the nAngptl4$^{WT}$ replicates. (C) Final simulation trajectory snapshots (t = 1000 ns) for the nAngptl4$^{E40K}$ replicates.

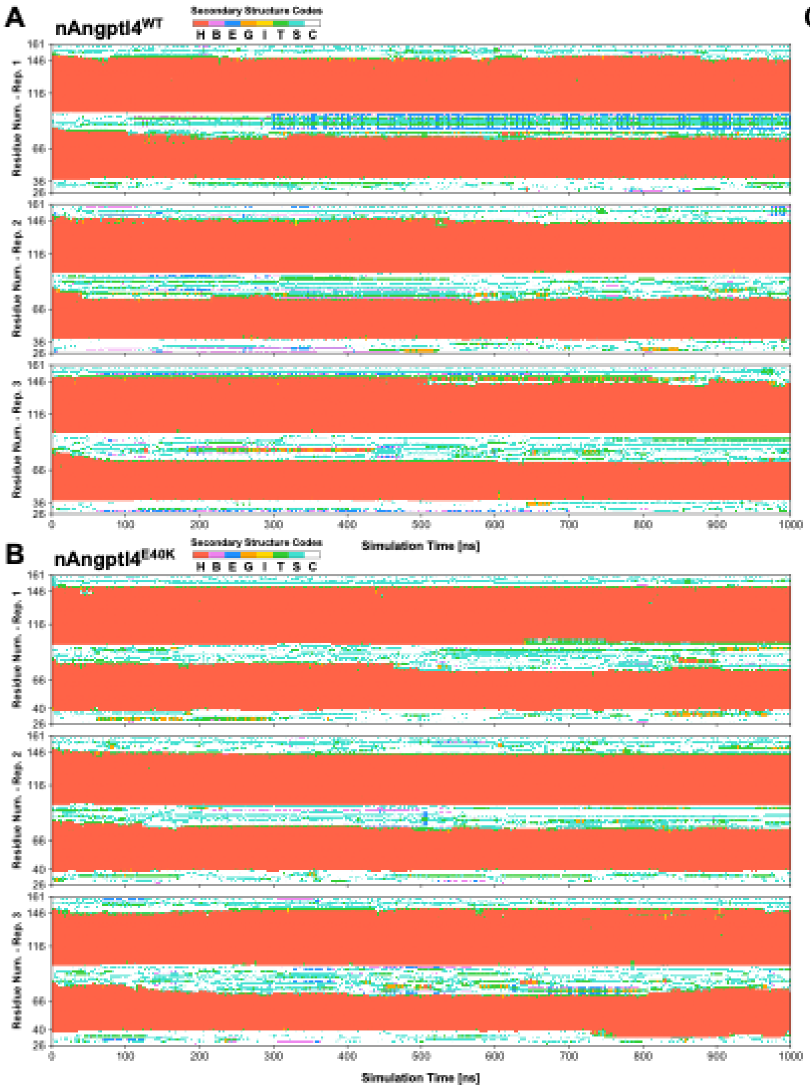


C

| Variant | α-Helicity Mean + SD |
|---|---|
| WT (n = 3) | 0.522 ± 0.013 |
| E40K (n = 3) | 0.543 ± 0.022 |
| Welch's t-test: t = -1.421, p = 0.249 | |

**Supplementary Figure 10.** (A) DSSP profiles for nAngptl4$^{WT}$ simulations over simulation time. Secondary structure codes include: α-helix (H), residue in isolated β-bridge (B), extended strand participating in β-ladder (E), 3/10-helix (G), π-helix (I), hydrogen-bonded turn (T), bend (S), and loop/irregular elements (C). (B) DSSP profiles for nAngptl4$^{E40K}$ simulations over simulation time. Secondary structure codes are the same as in (A). (C) Fractional α-helicity mean + standard deviation (SD) for WT and E40K, with a Welch's t-test performed across n=3 replicates per variant.

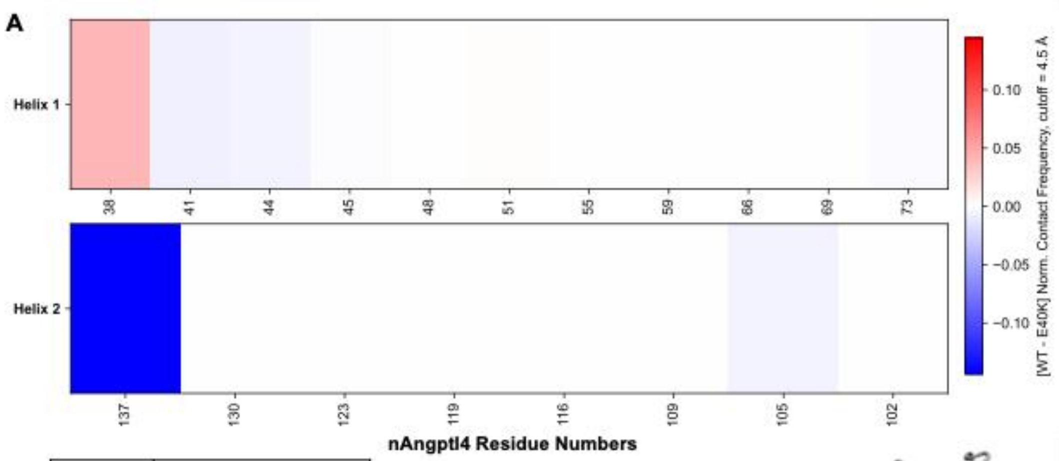


B

| Residue | Welch's t-test (n = 3) |
|---|---|
| 38 | t = 0.770, p = 0.485 |
| 41 | t = -1.890, p = 0.199 |
| 44 | t = -0.686, p = 0.553 |
| 73 | t = -1.000, p = 0.423 |
| 105 | t = -0.800, p = 0.487 |
| 137 | t = -1.060, p = 0.365 |

| Residue | Hedge's g test (n = 3) |
|---|---|
| 38 | g = 0.503 |
| 41 | g = -1.234 |
| 44 | g = -0.448 |
| 73 | g = -0.653 |
| 105 | g = -0.523 |
| 137 | g = -0.692 |

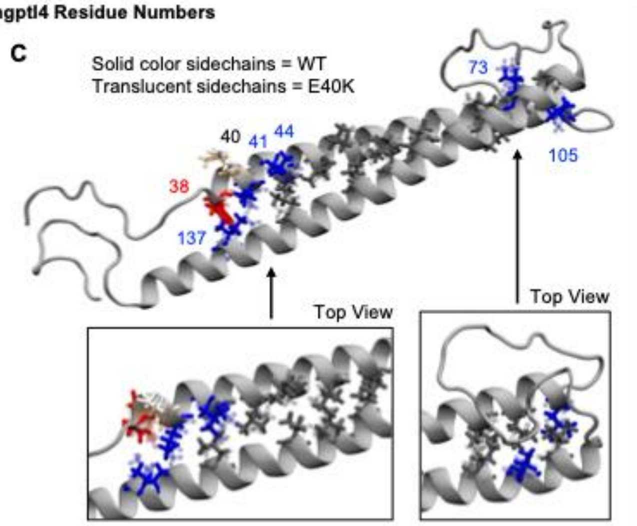


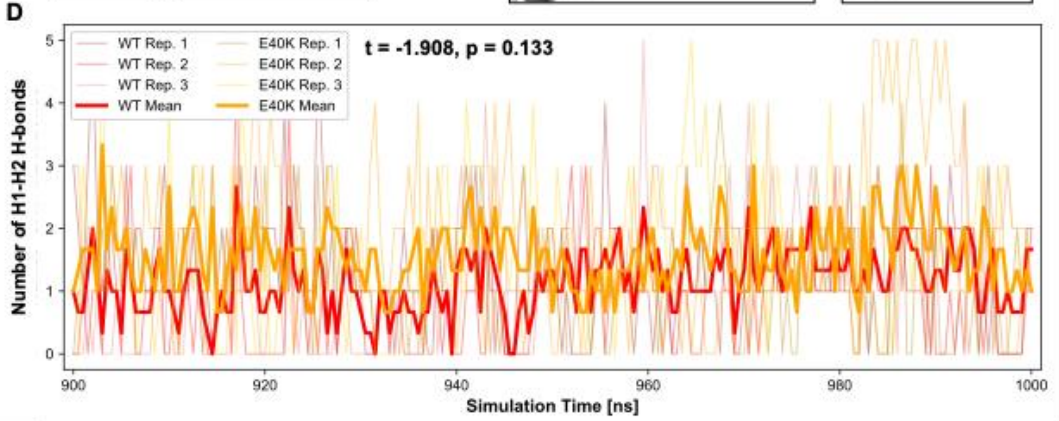


**Supplementary Figure 11**. (A) Differences in contact frequency across inward-facing, hydrophobic inter-helical residue contacts within nAngptl4 over the last 100 ns. Red values represent when WT has more contact. Blue values represent when E40K has more contact, comparatively. (B) Mean differences in contact frequency from WT to E40K were measured, with a Welch's t-test and Hedge's g test performed across n=3 replicates per variant. (C) Post-simulation representative structure of nAngptl4 with inward-facing, hydrophobic residues shown. Solid sidechains correspond to WT and translucent ones correspond to E40K. Residue 40 is in white for WT and orange for E40K. Differential contact residues are shown in red and blue, based on values in (A). Those with no differences are seen in dark gray. (D) Number of H-bonds between Helix 1 (H1) and Helix 2 (H2) over the last 100 ns of simulation time. Individual replicates are shown in faded colors, with WT in reds and E40K in oranges. The mean is shown in the bright, bolder lines (WT, red; E40K, orange). A Welch's t-test was performed for the mean across n=3 replicates per variant.

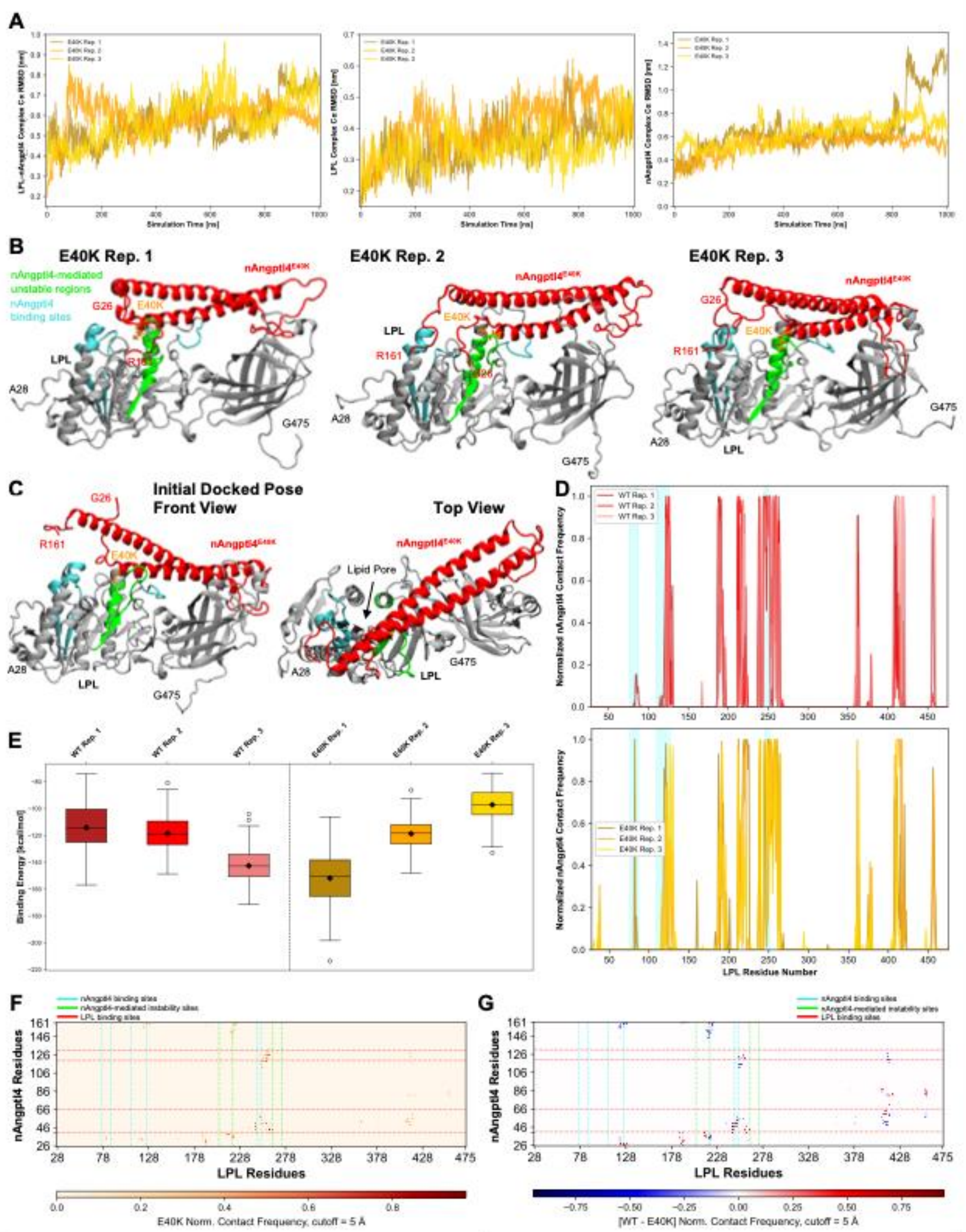


**Supplementary Figure 12.** (A) RMSD of Cα backbone of the LPL–nAngptl4$^{E40K}$ complex (left), LPL within the complex (center), and nAngptl4$^{E40K}$ within the complex (right) over simulation time. (B) Front view of LPL bound to nAngptl4$^{E40K}$ (final simulation time point, t = 1000 ns) for the three simulations. LPL is seen in gray, with the nAngptl4 binding sites in cyan (aa 76-87, 109-126, 245-251; LPL numbered 1-475) and allosterically unstable regions in green (aa 205-220, 264-274). nAngptl4$^{E40K}$ is seen in red and residue 40 in orange. (C) Initial docked pose for simulations with the front view (top) and top view (bottom). Protein colors are the same as described in (B). (D) Normalized contact frequency of nAngptl4$^{WT}$ (top) and nAngptl4$^{E40K}$ (bottom) with LPL, across LPL residues, from 900-1000 ns. Contact ranges from [0,1], with 0 and 1 representing no contact and always in contact, respectively. Cyan regions represent nAngptl4 binding sites on LPL. WT plot is identical to that in Fig. 2C. (E) Binding interaction energies ($\Delta G_{bind}$) between nAngptl4 and LPL from 900-1000 ns per simulation. The standard deviation, mean (diamond) and median (line) are shown. (F) Normalized (Norm.) contact map of nAngptl4$^{E40K}$ and LPL residues across the three simulations, from 900-1000 ns. Dashed lines represent bounds for binding sites on LPL and nAngptl4, for one another. (G) Differential normalized contact map of nAngptl4 interactions with LPL, from 900-1000 ns. Colored bounds are the same as in (F).

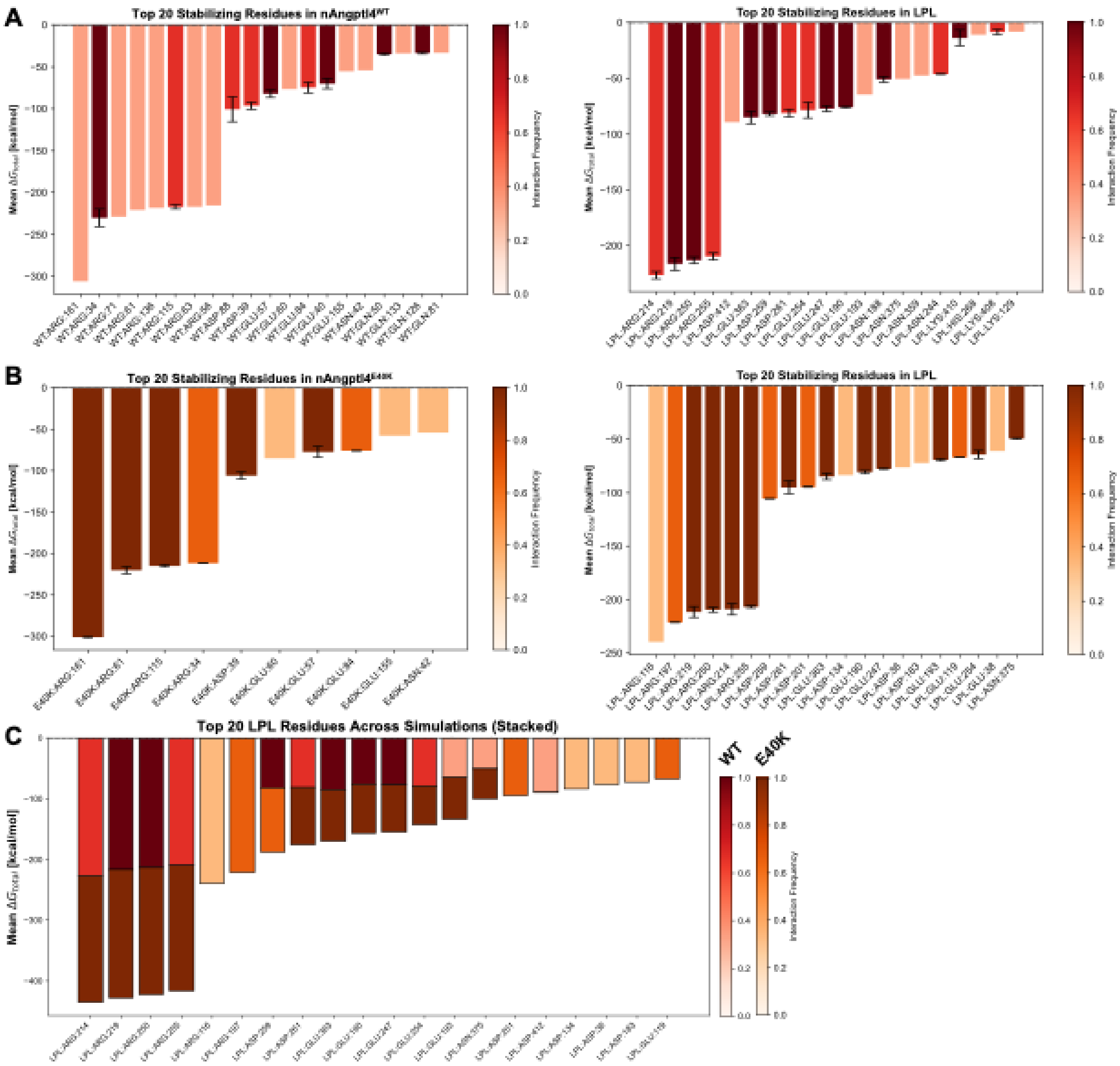


**Supplementary Figure 13.** (A) Summaries of binding energetic contributions across the top 10 H1 residues (left) and top 20 LPL residues (right). The mean $\Delta G_{bind}$ value across the replicates is shown with the standard deviation for residues involved in all 3 simulations. Colors correspond to the interaction frequency across the simulations, with scale bars shown on the right. (B) Summaries of binding energetic contributions across the top 10 H1-E40K residues (left) and top 20 LPL residues (right). The mean $\Delta G_{bind}$ value across the replicates is shown with the standard deviation for residues involved in all 3 simulations. Colors correspond to the interaction frequency across the simulations, with scale bars shown on the right. (C) Summaries of binding energetic contributions across the top 20 LPL residues in nAngptl4 interactions. The $\Delta G_{bind}$ value per nAngptl4 variant is shown, with the interaction frequency as the color. Interaction frequency scale bars are shown on the right, per variant.

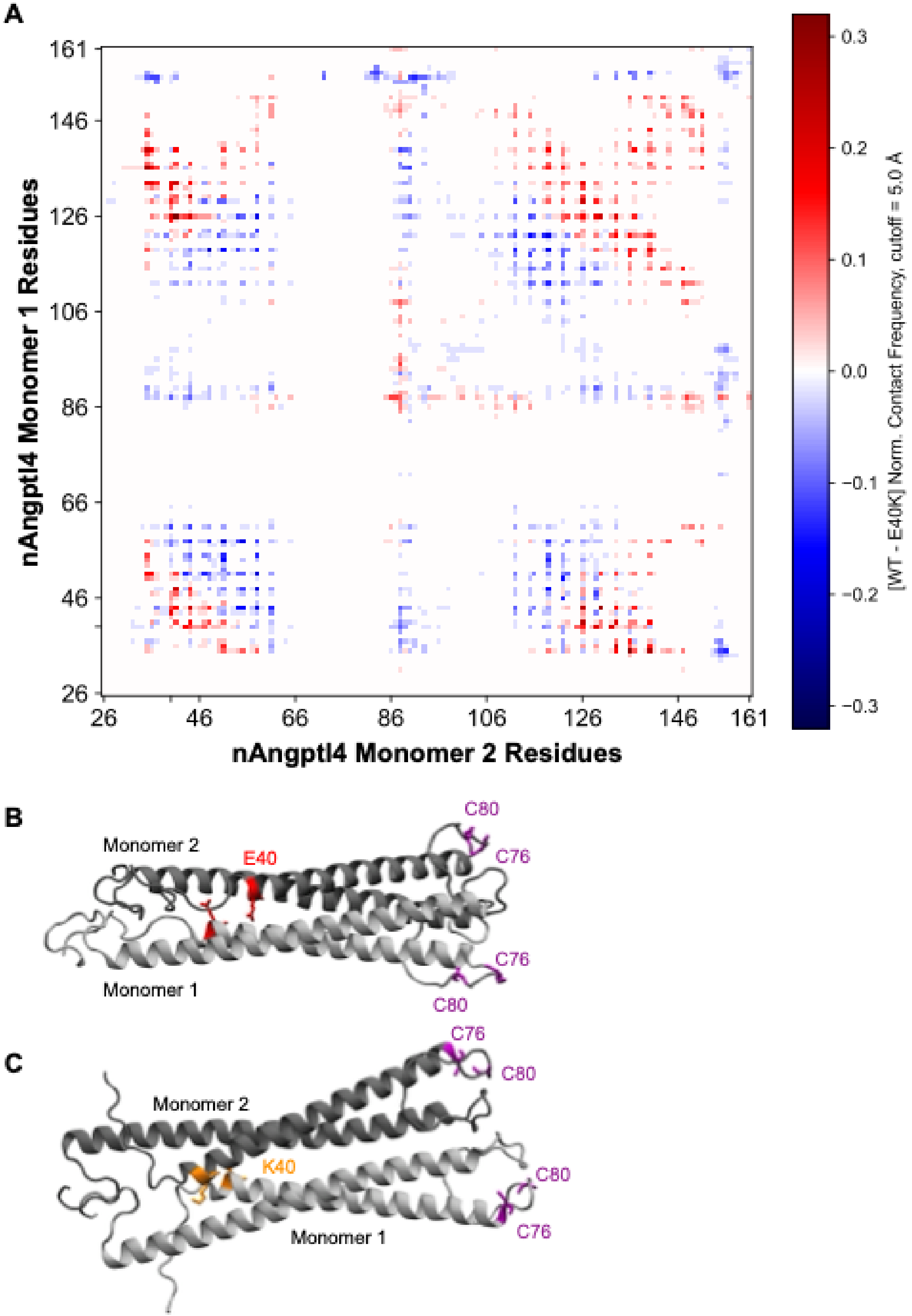


**Supplementary Figure 14**. (A) Differences in contact frequency across inter-residue contacts from one nAngptl4 monomer to another, in the top 50 docked, dimeric models (WT-WT or E40K-E40K). Red values represent when WT has more contact. Blue values represent when E40K has more contact, comparatively. (B) Top docked model of dimeric nAngptl4$^{WT}$ with residue 40 in red and hinge cysteines in purple. (C) Top docked model of dimeric nAngptl4$^{E40K}$ with residue 40 in orange and hinge cysteines in purple.

**Two-way repeated measures ANOVA with multiple comparisons with a Šidák correction**
Matching: Stacked
Sphericity assumed
α = 0.05

| Source of variation | % of total variation | P value | P value summary | Significant? |
|---|---|---|---|---|
| Time x nAngptl4 vs. LPL | 14.66 | <0.0001 | **** | Yes |
| Time | 37.63 | <0.0001 | **** | Yes |
| nAngptl4 vs. LPL | 44.6 | <0.0001 | **** | Yes |
| Subject | 2.176 | <0.0001 | **** | Yes |

| Components | Number |
|---|---|
| Columns (-/+ nAngptl4) | 2 |
| Time points | 13 |
| Subjects | 8 total (n=4, per column) |

**Supplementary Table 1. Two-way ANOVA with repeated measures was performed with multiple comparisons with a Šidák correction, for LPL-mediated hydrolysis of a fluorescent triglyceride analog with and without nAngptl4.**

**Two-way repeated measures ANOVA with multiple comparisons with a Tukey correction**
Matching: Stacked
Sphericity not assumed
α = 0.05

| Source of variation | % of total variation | P value | P value summary | Significant? |
|---|---|---|---|---|
| Time x Peptide | 2.242 | 0.0018 | ** | Yes |
| Time | 83.27 | <0.0001 | **** | Yes |
| Peptide | 10.19 | 0.0002 | *** | Yes |
| Subject | 2.65 | <0.0001 | **** | Yes |

| Components | Number |
|---|---|
| Columns (-/+ Peptide) | 4 |
| Time points | 13 |
| Subjects | 16 total (n=4, per column) |

| Tukey's multiple comparisons test, at t = 60 min | Mean diff. | Adjusted P value | P value summary | Significant? |
|---|---|---|---|---|
| H1 vs. LPL | 2490 | 0.0065 | ** | Yes |
| H1-E40K vs. LPL | 3964 | 0.018 | * | Yes |
| H2 vs. LPL | 2923 | <0.0001 | **** | Yes |

**Supplementary Table 2. Two-way ANOVA with repeated measures was performed with multiple comparisons with a Tukey correction, for LPL-mediated hydrolysis of a fluorescent triglyceride analog with and without nAngptl4-derived peptides.**

**Two-way repeated measures ANOVA with multiple comparisons with a Tukey correction**
Matching: Stacked
Sphericity not assumed
α = 0.05

| Source of variation | % of total variation | P value | P value summary | Significant? |
|---|---|---|---|---|
| Time x Peptides, nAngptl4 | 8.104 | <0.0001 | **** | Yes |
| Time | 43.71 | <0.0001 | **** | Yes |
| Peptide, nAngptl4 | 27.8 | 0.0061 | ** | Yes |
| Subject | 18.81 | <0.0001 | **** | Yes |

| Components | Number |
|---|---|
| Columns (-/+ Peptide, nAngptl4) | 5 |
| Time points | 13 |
| Subjects | 20 total (n=4, per column) |

| Tukey's multiple comparisons test, at t = 60 min | Mean diff. | Adjusted P value | P value summary | Significant? |
|---|---|---|---|---|
| H2 vs. LPL | -3525 | 0.0021 | ** | Yes |
| H2 vs. nAngptl4 | 1283 | 0.0087 | ** | Yes |
| H1-E40K vs. nAngptl4 | 2884 | 0.0185 | * | Yes |
| LPL vs. nAngptl4 | 4809 | 0.0013 | ** | Yes |

**Supplementary Table 3. Two-way ANOVA with repeated measures was performed with multiple comparisons with a Tukey correction, for LPL-mediated hydrolysis of a fluorescent triglyceride analog with and without nAngptl4-derived peptides, in the presence of nAngptl4.**
